# Functional traits drive speciation in tropical palms through complex interactions between genome size, adaptation and allometry

**DOI:** 10.1101/2025.09.07.673473

**Authors:** Sreetama Bhadra, Ilia J. Leitch, Sidonie Bellot, William J. Baker, Renske E. Onstein

## Abstract

- The importance of functional trait evolution and genome size on plant speciation are well established, but their interactive effects remain untested in a single comparative macroevolutionary framework.
- We integrated phylogenetic, trait and genome size data for palms (Arecaceae) – a large pantropical family (>2600 species) with 167-fold variation in trait and 60-fold variation in genome size. We used structural equation modelling to test three key hypotheses: trait evolution promotes speciation (H1: trait flexibility hypothesis), and, speciation and trait evolution rates are constrained by allometry (H2: allometric constraint hypothesis) and genome size (H3: large genome constraint hypothesis).
- We detected seven major speciation rate shifts during the ca. 110-million-year history of palms. Tip-derived speciation rates increased with faster evolution in leaves and plant height, supporting H1, whereas correlated evolution between trait evolution rates indirectly influenced speciation, supporting H2. Large genomes decreased plant height and stem diameter evolutionary rates, but increased leaf size evolution and speciation rates, thus partly supporting H3.
- Our findings illustrate how the complex interplay between genome size, allometry and trait evolvability affect speciation, emphasizing the importance of holistic approaches in macroevolution. Furthermore, our results point to potential general mechanisms driving speciation rates throughout the plant Tree of Life.

## Introduction

Speciation, the process by which new species arise, is the primary force shaping biodiversity. Speciation rates vary greatly across the Tree of Life (Reaney *et al*., 2020, Hernández-Hernández *et al*., 2021), yet why some lineages speciate more rapidly than others remains elusive. In plants, functional traits and genomic factors are well-established determinants of speciation, as they may respond to selective evolutionary forces (Ng & Smith, 2014, Pellicer *et al*., 2018, Onstein, 2020). However, how these factors interact to influence speciation rate variation across the plant Tree of Life is still poorly understood.

Functional trait evolution or ‘trait flexibility’ (Onstein, 2020) (i.e., the evolvability of traits through macroevolutionary time) influences growth, reproduction and survival of species, thereby playing a key role in adaptation, ecological speciation (Nosil, 2012), and the evolutionary radiation of lineages (‘trait flexibility hypothesis’) (Rolshausen *et al*., 2018, Nürk *et al*., 2020, Onstein, 2020). For example, the repeated innovation of key traits in angiosperms allowed them to colonize and diversify in new adaptive zones (*sensu* Simpson (Simpson, 1953)) and provided the functional machinery to rapidly adapt to environmental changes in the Cenozoic (last 66 million years) (Crepet & Niklas, 2009, Onstein, 2020). More recently, this pattern is exemplified by the association of angiosperm diversification with seed size (Igea *et al*., 2017, Onstein *et al*., 2017) and plant height (Boucher *et al*., 2017). While evolution of individual traits can open up new ecological opportunities, single traits rarely act in isolation to shape speciation.

Indeed, functional traits are structured in integrated phenotypes or trait syndromes, as a result of trait correlation – a pattern observed across diverse plant lineages (Niklas, 2004, Díaz *et al*., 2016). Consequently, changes in one trait, e.g., caused by genetic, developmental or evolutionary/adaptive change, can directly or indirectly influence other structurally or functionally-related traits. For instance, fruit size allometrically relates to stem diameter and overall plant architecture (Niklas, 2004), stem diameter growth tightly links to tree height (Rich *et al*., 1986, Zhao *et al*., 2021), and tree height and width often relates significantly with the diameter and lateral spread of roots across trees (Tumber-Dávila *et al*., 2022). Moreover, biomass allocation among roots, leaves and stems often follow predictable allometric rules (Niklas & Enquist, 2002). Such interdependencies, or allometric constraints, can limit the evolutionary flexibility of individual traits (‘the allometric constraint hypothesis’ (Harvey & Pagel, 1991)), thereby shaping trajectories of trait evolution and ultimately influencing speciation.

Functional trait evolution and speciation are further mediated by genomic factors. Genome size (total amount of DNA in a cell’s nucleus) biophysically constrains the minimum size of a cell (‘the nucleotypic effect’ (Bennett, 1971)), which affects anatomical (e.g., guard cell size, cell packing density) and physiological (e.g., stomatal conductance, maximum photosynthetic rate) traits (Roddy *et al*., 2020). The nucleotypic effect of genome size also influences evolution of morphological traits like seed size (Lopes *et al*., 2021, Carta *et al*., 2022) and pollen size (Glazier, 2021), and thus potentially the overall adaptability and speciation of plants (‘large genome constraint hypothesis’ (Knight *et al*., 2005)) (Bhadra *et al*., 2023). For example, reductions in genome size (genome downsizing) of angiosperms has been linked to their global speciation and rise to dominance during the Cretaceous (Simonin & Roddy, 2018).

Although macroevolutionary associations between traits and speciation (Ng & Smith, 2014, Onstein, 2020), genome size and speciation (Pellicer *et al*., 2018, Moeglein *et al*., 2020), genome size and traits (Roddy *et al*., 2020, Glazier, 2021, Lopes *et al*., 2021, Carta *et al*., 2022), and functional traits and allometry (Rich *et al*., 1986, Lauri, 2019) are well established, their interactive effects on speciation remain poorly understood. To capture these multidimensional relationships, we moved beyond the traditional univariate approaches that test the temporal shifts in individual trait evolution which potentially overlook the complex interdependencies among these processes. Instead, we employed a multivariate structural equation modelling (SEM) framework integrating genome size with tip-derived estimates of trait evolution and speciation rates (Fig. 1). This enabled us to test three key hypotheses within a single analytical model (Fig. 1): first, that faster trait evolution leads to faster speciation (‘trait flexibility hypothesis’, H1); second, that trait evolution is constrained by allometry, thereby reducing speciation rates (the ‘allometric constraint hypothesis’, H2); and third, that large genomes constrain trait evolution and speciation rates (Knight *et al*., 2005) (‘large genome constraint hypothesis’, H3) – while capturing both direct and indirect effects.

**Fig. 1.**
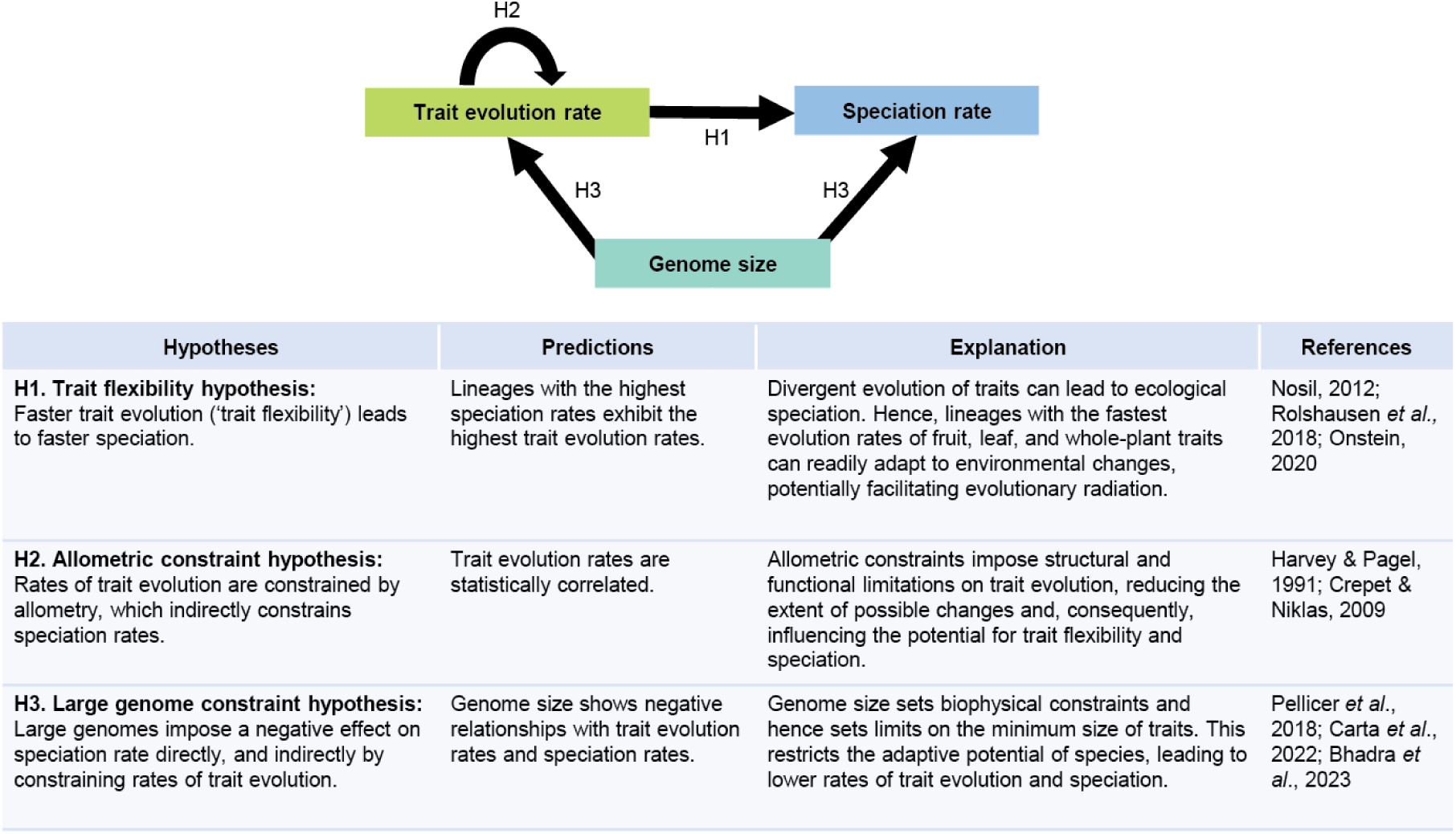
Conceptual framework of hypotheses and predictions on the impact of trait evolution rates and genome size on speciation rates. The diagram illustrates three hypothesized pathways (H1–H3) linking trait evolution rates, genome size, and speciation rates. H1 represents the ‘trait flexibility hypothesis’ which states that faster trait evolution promotes speciation. H2 represents the ‘allometric constraint hypothesis’ which states that allometric relationships between trait evolution rates impose evolutionary constraints that indirectly influence speciation. H3 represents the ‘large genome constraint hypothesis’ which states that genome size limits trait evolution rates and speciation rates. The table summarizes each hypothesis with its corresponding predictions, offering a mechanistic perspective on how these factors may interact to shape speciation dynamics. This framework is applied here to the palm family (Arecaceae).

We tested this framework on palms (Arecaceae), a species-rich tropical plant clade comprising ∼2,600 species (Govaerts *et al*., 2021), often dominating in rainforest floras (Kissling *et al*., 2019, Henderson, 2002). Anatomical constraints restrict palms largely to subtropical and tropical areas (Tomlinson, 2006, Reichgelt *et al*., 2018), reflecting strong niche conservatism. Despite this, palms have persisted through ca. 110-million-years of climatic changes (Baker & Couvreur, 2013), exhibiting remarkable adaptation and extraordinary trait variations. For example, they exhibit 167-fold variation in fruit/seed size (0.3 cm long in *Geonoma interrupta* to 50 cm long *Lodoicea maldivica*, the largest seed among angiosperms), and leaf blade length (0.15 m in *Hydriastele montana* up to 25 m in *Raphia regalis*, the largest leaf among angiosperms) (Tomlinson, 2006, Kissling *et al*., 2019). They also exhibit about 60-fold variation in genome size, largely independent of polyploidy and chromosome number variations (Pellicer & Leitch, 2020, Schley *et al*., 2022). This variation in functional traits and genome size makes palms an ideal system to understand how these factors may have facilitated or constrained palm speciation, resulting in their modern diversity.

## Materials and Methods

### Palm phylogeny

The palm phylogenetic tree by Faurby *et al*. (2016) that we used for our analyses is a time-calibrated Bayesian supertree that includes 184 genera and 2,528 species (∼100% species coverage) generated by using a mix of species-level genetic and morphological datasets (Faurby *et al*., 2016). This study includes MCC (Maximum Clade Credibility) tree which we used for the main analyses, and 1000 constrained, posterior-distribution trees, 100 of which were used for sensitivity analysis.

### Functional trait data

Species-level data for functional traits of palms were obtained from the PalmTraits v.1.0 database (Kissling *et al*., 2019) and the trait values were used with no further modification, except in cases where a normalization of the values was needed. We initially selected 11 quantitative traits characterizing the size of fruits (average, maximum and minimum length and width), leaves (maximum lengths of leaf blade, rachis and petiole), stem (maximum stem diameter) and plant height.

Fruit size and leaf size were represented by multiple categories in the database (Kissling *et al*., 2019). While the subsequent analyses using structural equation modeling (SEM; explained later) allows the testing of multiple variables within a single model, increasing the number of variables greatly raises the number of possible path topologies. This increases the intrinsic dimensionality of the dataset and can introduce type I errors (false positives) into the resulting model (Matthews *et al*., 2023). To reduce such errors and avoid the risk of mistaking random statistical patterns for biological relevance, it was essential to include only variables that contribute meaningful value to the model. To identify the relevant variables for fruit size and leaf size, we performed linear regression analyses in R to explore the relationships between the trait categories within these plant parts. The categories of fruit size and leaf size were strongly positively correlated (Fig. S1).

Hence, the following four traits, with the largest available dataset (Table S1), were chosen for further analysis: fruit size (measured as the average length of fruit, 80.23% species), leaf size (measured as the maximum length of leaf blade, 74.37% species), stem diameter (measured as the maximum diameter of stem, 75.82% species) and plant height (measured as the maximum whole plant height including the crown, 78.03% species).

### Genome size data

Genome size data (1C-values, in Gbp) were compiled from the Plant DNA C-values database (Pellicer & Leitch, 2020) and those published in Schley *et al*. (2022) (Table S1, S2), resulting in a dataset covering 16.71% of palm species. Four polyploid palms - *Arenga caudata* (2n=4x=64), *Jubaeopsis caffra* (2n=12x=160-200), *Rhapis humilis* (2n=4x=72) and *Voanioala gerardii* (2n=38x=596) were considered as outliers because all other palm species are diploids (Schley *et al*., 2022). These species were retained in the dataset but their impact was carefully monitored throughout sensitivity analyses (explained later).

### Taxonomic standardization

Most of our analyses were based on the phylogenetic tree (Faurby *et al*., 2016). Hence, we standardized species names across all data sources using the taxonomy in Faurby *et al*. (2016). To ensure clarity, we verified the taxonomy using the World Checklist of Vascular Plants (WCVP) (Govaerts *et al*., 2021) and recorded the accepted names (Table S2), and removed records that could not be unambiguously assigned to accepted species.

### Estimation of speciation rates

We estimated the present-day speciation rates of each species (= tip rates; measured as lineages per million years) using the MCC phylogenetic tree (Faurby *et al*., 2016) which includes ∼100% species utilizing three approaches since consensus on a single best method is lacking. This included two model-based approaches - BAMM (Rabosky, 2014) and ClaDS (Maliet & Morlon, 2022) - and one model-free approach - DR (Jetz *et al*., 2012). Each method differs in their assumptions to deduce speciation trajectories. Hence, the results of the present-day speciation rates are unlikely to be mutually consistent if the rate estimates are not firmly grounded on the data. Using Pearson’s *r* correlation, we tested the comparability of these three methods used for speciation rate estimates (Fig. S2).

BAMM uses Bayesian statistics to estimate discrete shifts in speciation and diversification rates at nodes of a phylogenetic tree. The *speciation-extinction* parameter simulates posterior distribution based on the prior rate shifts utilizing reversible-jump Markov Chain Monte Carlo that implements a compound Poisson process. This allows for rate heterogeneity and random shift distribution along the tree and through evolutionary time (Rabosky, 2014). Priors were obtained by the *setBAMMpriors* function in the ‘BAMMtools’ package in R (Rabosky *et al*., 2014) (Table S3) which resulted in *expectedNumberOfShifts* parameter to be 1. To check the sensitivity of the assumed priors on the estimated posterior distribution of BAMM speciation rates (Moore *et al*., 2016, Meyer *et al*., 2018), we compared speciation rates deduced from different prior values (see Notes S1, Fig. S3) and retained *expectedNumberOfShifts* parameter as 1. MCMC simulation consisted of four independent chains of 300 million generations each, with shift configuration being sampled every 10,000 steps. Run convergence was assessed by the *effectiveSize()* function of the package ‘coda’ (Plummer *et al*., 2006). The initial 10% of MCMC was discarded as burn-in (number of analyzed posterior samples=27,001) and BAMM output was analyzed using BAMMtools (Rabosky *et al*., 2014). Tip rates of speciation were extracted (Table S2) using the *getTipRates()* function and lineage-specific speciation were visualized as phylorate plots by *plot.bammdata()* function (Rabosky *et al*., 2014).

ClaDS uses a Bayesian approach to deduce speciation rate allowing for small and frequent rate shifts at each branching event of a phylogenetic tree (Maliet *et al*., 2019). We used the Julia language (Bezanson *et al*., 2017) implementation of the ClaDS model (ClaDS2) in the PANDA package (Morlon, 2024) that uses a constant turnover of extinction rates. The model uses data augmentation with three MCMC chains (Maliet & Morlon, 2022). Convergence was assessed using Gelman statistics (Gelman *et al*., 2013) and the chains stopped when the value dropped below 1.05 (Maliet & Morlon, 2022). To view the speciation rates superimposed on the palm phylogeny, the function *plot_ClaDS_phylo()* of the Rpanda package in R was used, and the tip rates of each species were extracted (Table S2).

DR is a model-free approach that estimates the present-day speciation rate of each species in a phylogeny by making minimal assumptions about the speciation process. It calculates tip rates as a weighted mean of inverse branch lengths under a time-constant, homogeneous model in the absence of extinction (Jetz *et al*., 2012). We used the ‘epm’ package (Jetz *et al*., 2012) in R to obtain the tip rates for each species (Table S2).

### Estimation of functional trait evolution rates

We implemented the *phenotypic evolution* parameter in BAMM v. 2.5.0 (Rabosky, 2014) to estimate the present-day rates of trait evolution (= tip rates of traits; measured as changes per million years) from the trait data of average fruit length (fruit size), maximum leaf blade length (leaf size), maximum stem diameter and maximum plant height with the MCC tree (Faurby *et al*., 2016) as the phylogenetic backbone. The tree was pruned to obtain sub-trees for each trait since data availability varied (Table S1) and the BAMM *phenotypic evolution* model does not implement sampling fractions (Igea *et al*., 2017, Beltrán *et al*., 2021). The *phenotypic evolution* parameter is based on a similar hypothesis as the *speciation-extinction* parameter (for details, see Section ‘*Estimation of speciation rates*’ above). We obtained the priors using the *setBAMMpriors* function of BAMMtools (Rabosky *et al*., 2014) (Table S2) that set the *expectedNumberOfShifts* to 1. Similar to BAMM speciation rates, sensitivity of assumed priors on the estimated posterior distribution of BAMM trait evolution rates were compared for different prior values for each trait (see Notes S1), and was retained as 1. MCMC simulations were run for 300 million generations for each of the four independent chains and the convergence of the runs were assessed by the ‘coda’ package (Plummer *et al*., 2006). The first 10% of results were discarded as burn-in (number of analyzed posterior samples=27,001). BAMM outputs were analyzed to get the tip rates of each species for each trait. Tip rates were extracted (Table S2) using the *getTipRates()* function and the lineage-specific trait evolution rates for each trait were visualized as phylorate plots using BAMMtools (Rabosky *et al*., 2014).

### Structural equation models

We used structural equation modeling (SEM) to disentangle the direct effects of genome size and functional traits evolution rates on the speciation of palms, of genome size on trait evolution rates, and among trait evolution rates. All variables were log-transformed and scaled between 0 and 1 to approach normality in model residuals, and to be able to compare standardized effect sizes. SEM was implemented using the ‘lavaan’ R package (Rosseel, 2012). Since the ‘lavaan’ package does not consider phylogenetic structure, we performed phylogenetic generalized least squares (PGLS) analysis on the final SEM model *a posteriori* (see next Section). We started with an *a priori* model that included all the hypothesized pathways between the variables (Fig. S4) based on pairwise correlations and theoretical considerations. Specifically, we expected that the evolution rates of all functional traits would positively influence speciation rates. In addition, the evolution rate of plant height was expected to positively influence stem diameter, leaf size and fruit size evolution. Similarly, stem diameter was expected to have a positive influence on leaf size and fruit size evolution, and leaf size was expected to positively impact fruit size evolution. Genome size was expected to impose a negative impact on speciation rate and on all trait evolution rates.

Starting with the *a priori* model, statistically insignificant relationships were progressively removed until each model only consisted of significant pathways (*p* < 0.05), resulting in the final model. We extracted the standardized coefficients of the final model. The modification indices were checked using the function *modindices()* to assess if any previously omitted parameters or covariates were required to improve model fit. Model fit was assessed using multiple indices-*p*-value of χ^2^ – tests >0.05, Comparative Fit Index (CFI) >0.95, Tucker-Lewis Index (TLI) >0.90, Root Mean Square Error of Approximation (RMSEA) <0.05, and Standardized Root Mean Square Residual (SRMR) <0.08. Three separate SEM analyses were conducted with speciation rates estimated from BAMM, ClaDS or DR. Most associations between the variables were consistently supported in all models.

### Phylogenetic autocorrelation

Hence, we evaluated whether main predictors of speciation as detected in the SEMs were also supported when accounting for phylogenetic dependence on the response variable (i.e., speciation rate) using phylogenetic generalized least squares (PGLS) as implemented by the R caper v 1.0.3 package (Orme *et al*., 2013) in R. Phylogenetic autocorrelation is accounted for in ‘caper’ by including a variance-covariance term into the model errors, derived from the shared branch lengths between species and scaled by the λ parameter (Freckleton *et al*., 2002). λ quantifies the strength of phylogenetic autocorrelation, where λ=0 indicates no phylogenetic signal where trait evolution is considered to be independent of their phylogenetic relationships, while λ=1 indicates trait evolution proportional to phylogenetic relatedness under Brownian motion. We estimated the λ parameter in the PGLS models using maximum likelihood, while fixing the other two scaling parameters (i.e., delta and kappa) to one.

### Phylogenetic tree topology and SEM robustness

A MCC tree is a consensus tree that best represents the overall phylogenetic signal in the data and does not account for the uncertainty of phylogenetic tree topology. This may have an impact on the estimates of the correlations of the SEM model since topological changes can influence speciation rates which can potentially affect the estimates deduced by SEM. To account for the uncertain topology of the palm phylogeny, we deduced speciation rates from 100 randomly sampled constrained, posterior-distribution trees (Faurby *et al*., 2016) using BAMM.

The robustness of the final model estimates was tested by comparing this model with the range of estimates (standardized coefficients) obtained by reiterating the final SEM model 100 times using the variable set of speciation rates derived from BAMM. Coefficients of each correlation were extracted and their distribution was plotted to check for variation while locating the estimate of MCC SEM on the plot. We also checked the significance of *p*-values for each of the analyses.

### Effect of large genome sizes and polyploidy

We repeated the SEM analysis using BAMM speciation rates after removing the four polyploid species from our dataset to test the effect of outliers. To test the influence of large genomes on speciation and trait evolution rates, we repeated SEM using BAMM speciation rates with two subsets of genome size data: species with genome sizes larger than the median value (i.e., genome sizes larger than 2.64 Gbp/1C) and species with genome size in the 75th percentile range (i.e., genome sizes above the 75th percentile threshold of 4.26 Gbp/1C). In all cases, we started with the *a priori* model (Fig. S4) and progressively removed the insignificant relations until the model possessed only significant pathways, and the fit indices had required values.

## Results

### Palm speciation is associated with high rates of trait evolution

At least seven evolutionary radiations were identified in palms, characterized by lineages with speciation rates (λ_Sp_= 0.234-0.358 lineages/myr; range reflects variation across radiations in BAMM) exceeding the background speciation rate in palms (average λ_Sp_= 0.178-0.260 lineages/myr; range reflects estimates across 100 phylogenetic trees in BAMM) (for details of all speciation rates, see Table S2). The radiations occurred in three of the four non-monotypic subfamilies, specifically in tribe Calameae (subtribe Calaminae) of subfamily Calamoideae, tribes Cryosophileae and Trachycarpeae (subtribe Livistoninae) of subfamily Coryphoideae, and tribes Cocoseae (subtribe Attaleinae) and Areceae (subtribes Dypsidinae, Ptychospermatinae and Arecinae) of subfamily Arecoideae, which together included 24 out of the 184 palm genera currently recognized (Figs. 2, S5, Table S2).

**Fig. 2.**
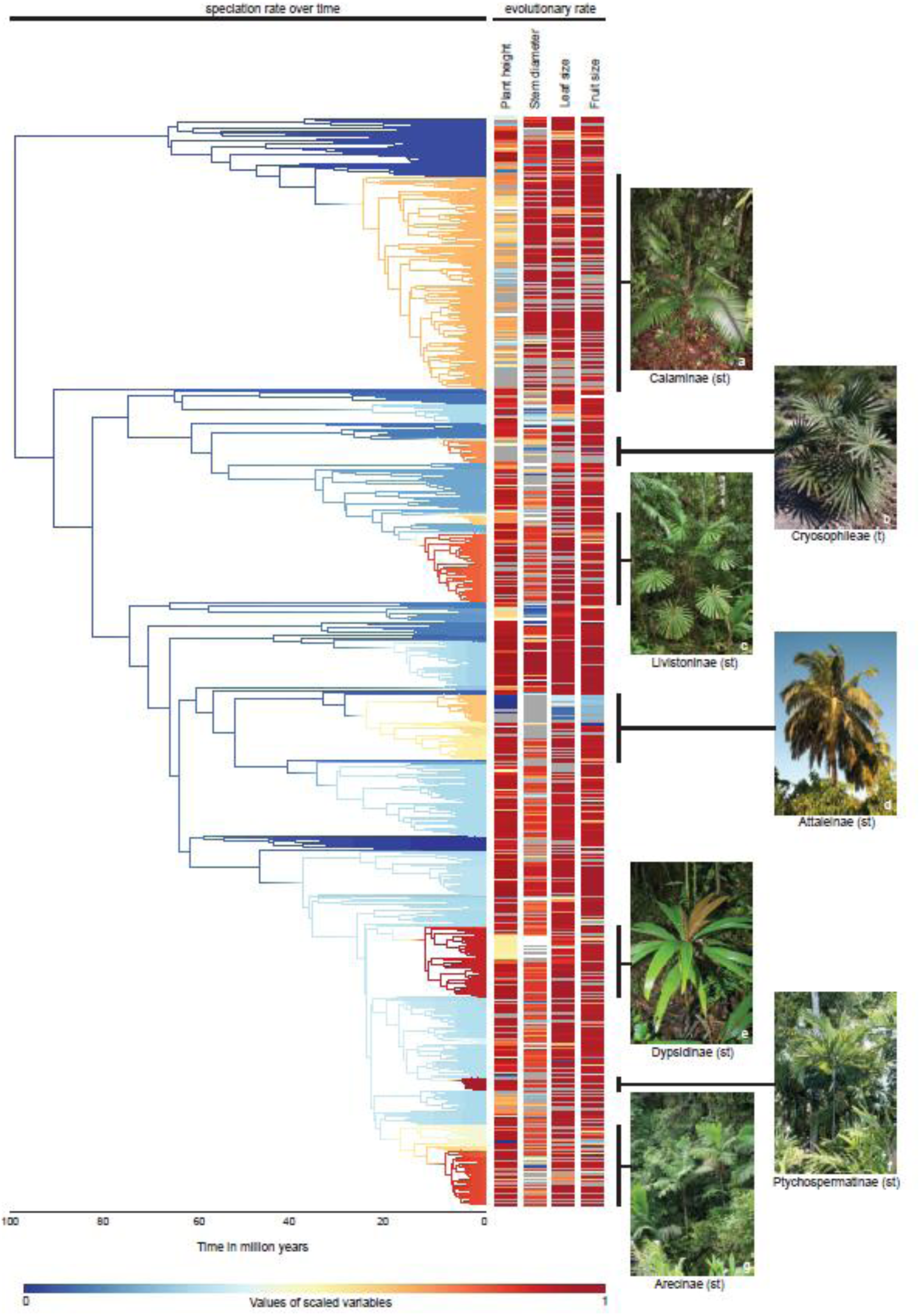
Evolutionary radiations and shifts in speciation rates in palms (Arecaceae). The phylogeny of 2,528 palm species with associated data on evolutionary rates of plant height, stem diameter, leaf size and fruit size (for details on species names, see Fig. S5). The phylogeny is colour-coded with speciation rates estimated using BAMM (Bayesian Analysis of Macroevolutionary Mixtures). Evolutionary rates of each trait per species, also estimated by BAMM, are indicated alongside the tree. Grey denotes missing data. Vertical black bars on the right highlight lineages exhibiting evolutionary radiations (i.e., speciation rates estimated to be above background speciation rates). Names of these lineages (t: tribes; st: subtribes) are shown below each figure featuring a representative species: (a) *Calamus korthalsii* (b) *Coccothrinax miraguama* (c) *Licuala lauterbachii* (d) *Cocos nucifera* (e) *Dypsis mocquerysiana* (f) *Ptychosperma keiense* (g) *Pinanga rumphiana*. Photos by William J. Baker.

Structural equation modeling tested the relationship between trait evolution rates (regardless of whether trait values increased or decreased) and genome size with palm tip-derived speciation rates. Our final SEM model (*N*=372 species, Fig. 3) integrates findings derived from speciation rate estimates from BAMM, ClaDS and DR, which produced largely consistent patterns (Figs. S6-S9). Our final model indicates that our combined predictors explain up to 12% variation in speciation rates and up to 19% in trait evolution rate variation across species (Fig. 3). Rates of leaf size (standardized coefficient (std. coeff.) = 0.151; *p*-value<0.05), and plant height (std. coeff. = 0.098; *p*<0.05) evolution are positively associated with speciation rates, in support of the trait flexibility hypothesis (H1). In contrast, the rate of stem diameter evolution was negatively associated with the speciation rate (std. coeff. = −0.159; *p*<0.05). However, it is important to note that the stem diameter evolution-speciation association was only supported by SEMs using BAMM and ClaDS (Figs. S6, S7), and the plant height evolution-speciation link was only supported in SEMs with ClaDS and DR (dashed lines in Fig. 3; Fig. S7, S8). Fruit size evolution showed no direct relationship with speciation. Phylogenetic autocorrelation analyses of the main predictors of speciation rates indicted that stem diameter evolution and speciation rates were phylogenetically correlated, and therefore these relationships should be interpreted with caution (Table S4).

**Fig. 3.**
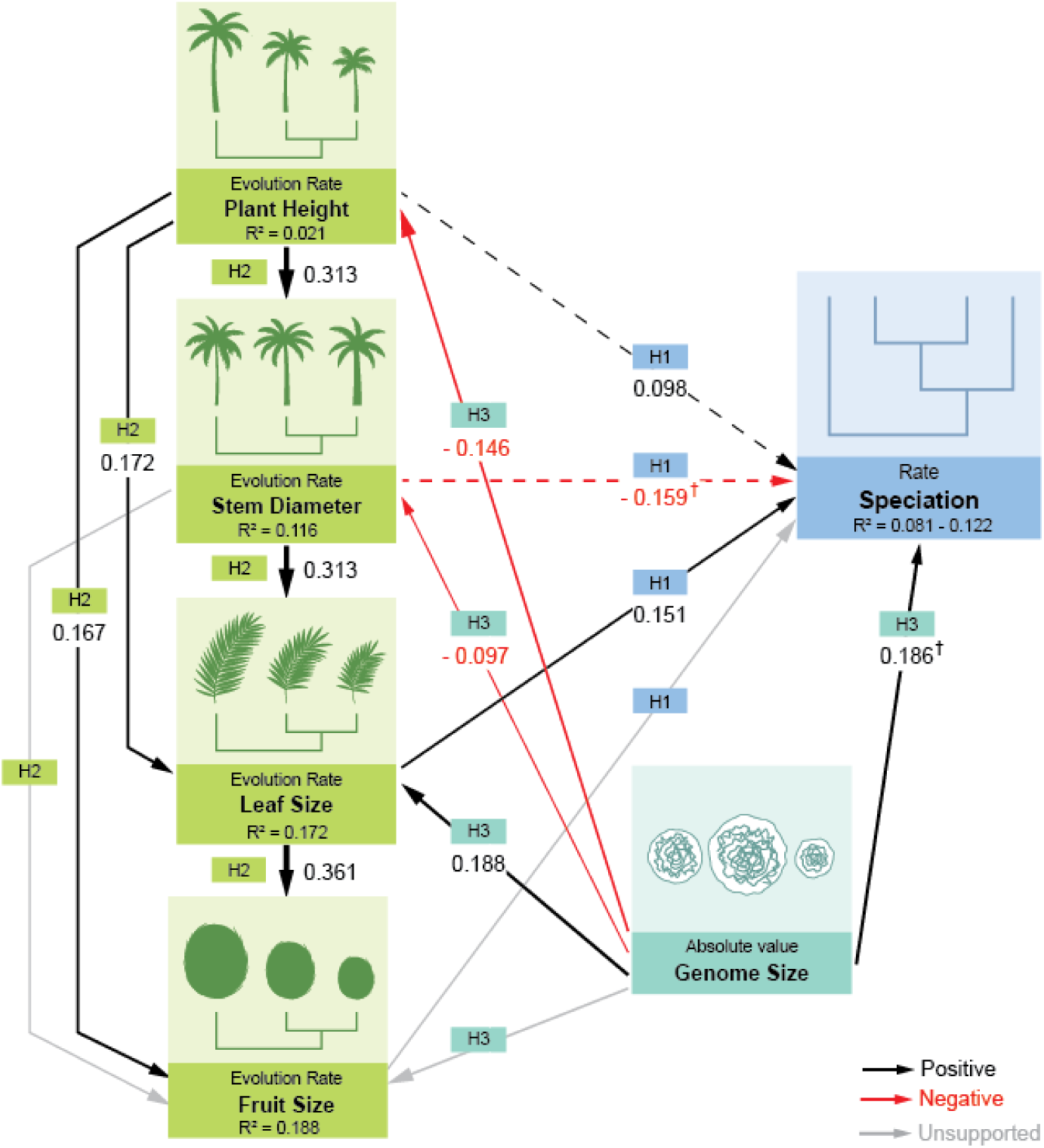
Trait flexibility, allometric constraints and genome size effects on speciation rates in palms (Arecaceae). Structural equation model (SEM) showing the standardized effects of trait evolutionary rates of plant height, stem diameter, leaf size and fruit size on speciation rates (trait flexibility hypothesis, H1), correlations between trait evolution rates (allometric constraint hypothesis, H2), and of genome size on speciation (genome size constraint hypothesis, H3) in palms (*N*=372 species). Speciation rates were derived from BAMM (Bayesian Analysis of Macroevolutionary Mixtures), ClaDS (Cladogenetic Diversification rate Shift) and DR (Diversification Rate statistics). The effect sizes indicate standardized coefficients with significance (p<0.05). The arrow thickness is proportional to coefficient values and the arrow direction represents the direction of effects. Black arrows denote positive effects and red arrows denote negative effects. Dotted arrows represent effects that were statistically supported in only one or two of the speciation rate scenarios derived from BAMM, ClaDS and DR (for details, see Figs. S6-S8). Coefficient values of BAMM are indicated, or from the respective models if supported only in ClaDS or DR. Grey arrows denote tested but statistically unsupported effects in any of the scanerios (p>0.05). The R^2^ values (explained variation based on the combined predictors) are consistent across models (BAMM, ClaDS, DR) for all variables, except for speciation rates, hence the reported range. Model fit indices confirmed an adequate fit of all models: *p*-value of *χ*2 tests >0.05, Comparative Fit Index (CFI) >0.95, Tucker-Lewis Index (TLI) >0.90, Root Mean Square Error of Approximation (RMSEA) <0.05, and Standardized Root Mean Square Residual (SRMR) <0.08. † indicates relationships not supported in any of the phylogenetic generalized least squares (PGLS) analyses (for either BAMM, ClaDS or DR-derived speciation rates; for details, see Table S4).

### Trait evolution is constrained by allometry

In support of the allometric constraint hypothesis (H2), our SEM indicated positive correlations among trait evolution rates (Figs. 3, S6-S8). Specifically, increases in rates of leaf size (std. coeff.= 0.361; *p*<0.05) and plant height (std. coeff.= 0.167; *p*<0.05) evolution were associated with increases in the rate of fruit size evolution. Similarly, faster rates of evolution of plant height (std. coeff.= 0.172; *p*<0.05) and stem diameter (std. coeff.= 0.313; *p*<0.05) were associated with an increased rate of leaf size evolution. A positive association was also found between rates of plant height and stem diameter evolution (std. coeff.= 0.313; *p*<0.05). Thus, through their allometric relationship with leaf size, plant height and stem diameter evolution may indirectly influence speciation rate.

### Genome size constrains trait evolution and, hence, speciation

We found that large genomes were positively associated with speciation rates (std. coeff.= 0.186; *p*<0.05) (Fig. 3). This contradicts the expectations from the ‘large genome constraint hypothesis’ (Knight *et al*., 2005), and rejects H3. This relationship persisted after reanalyzing the SEMs excluding four polyploid species to account for the potential impacts of recent polyploidization on speciation (Fig. S10), and when analyzing sub-sets of species with above median genome sizes (i.e., genome sizes larger than 2.64 Gbp/1C, *N*=189 species, Fig. S11) and those above the 75^th^ percentile (i.e., genome sizes larger than 4.26 Gbp/1C, *N*=95 species, Fig. S12). However, the PGLS analyses for BAMM-, ClaDS- and DR-derived speciation rates all indicated that the effect of genome size on speciation rates was no longer significant when correcting for phylogenetic autocorrelation (Table S4).

In terms of trait evolution, genome size was negatively associated with rates of plant height (std. coeff.= −0.146; *p*<0.05) and stem diameter (std. coeff.= −0.097; *p*<0.05) evolution, but positively associated with the rate of leaf size evolution (std. coeff.= 0.188; *p*<0.05). Together, these results suggest that genome size may influence palm speciation rates indirectly, through its effects on functional trait evolution, thus partially supporting the ‘large genome constraint hypothesis’ (H3).

## Discussion

Our study highlights that palm speciation is shaped by a complex interplay of trait evolution, allometric relationships and genome size. Using an integrative macroevolutionary approach, we identified potential direct and indirect pathways influencing palm speciation, offering a comprehensive framework to understand the mechanisms behind the extraordinary diversity of this tropical plant family. Although morphological divergence during allopatric and sympatric speciation may be rare when comparing sister species (Olivares *et al*., 2025), our results suggest that shifts in trait evolution rates as detected among larger clades may closely link to speciation rate shifts, embellishing palms as a potential model for studying ecological speciation in plants. The strong correlations revealed by our study do not inherently establish causation, but our findings are grounded in a robust body of evidence that supports causal connections.

### Trait flexibility shapes palm speciation through divergent morphological evolution

Speciation across the plant Tree of Life has often been linked to morphological evolution (Crepet & Niklas, 2009, Boucher *et al*., 2017, Igea *et al*., 2017, Onstein, 2020), and our results suggest that palms follow a similar pattern. We found that rapid rates of leaf size and plant height evolution are associated with higher speciation rates, supporting the ‘trait flexibility hypothesis’ (H1) (Figs. 3, S6-S8). For example, genera such as *Pinanga* and *Ptychosperma* (tribe Areceae) exhibit high rates of leaf size and plant height evolution alongside fast speciation (Fig. S5, Table S2). These results suggest that divergent trait evolution (‘flexibility’) over macroevolutionary timescales can promote ecological speciation, perhaps through enhanced adaptive potential. Leaf size evolution, coupled with variations in leaf shape (Torres Jiménez *et al*., 2023), can optimize temperature regulation and light capture under diverse environmental conditions (Wright *et al*., 2017). For example, small-leaved palms of tribe Cocoseae, such as species within *Bactris,* dominate the warm and humid central Amazon, whereas their large-leaved relatives, such as *Attalea*, are more prevalent in the drier northwestern and southeastern Amazon (Göldel *et al*., 2015). The ability to adjust leaf size can also improve carbon gain efficiency enhancing adaptability to habitats of different light intensity, such as of understorey palms (e.g., *Pinanga coronata*) to shaded habitats (Ma *et al*., 2015). Plant height evolution offers biomechanical support to foliar architecture in tall, solitary-stemmed palms, enabling canopy access to light critical for photosynthesis in forest environments (Göldel *et al*., 2015). This trait-driven partitioning of vertical space (understorey vs. canopy) can reduce interspecific competition and facilitate ecological speciation through spatial isolation (Poorter & Rozendaal, 2008, Li *et al*., 2020). Additionally, vertical stratification can affect seed dispersal dynamics by segregating frugivore communities along vertical gradients (Onstein *et al*., 2017). This is further supported by the positive association between plant height and fruit size evolution rates in our study. Small-bodied understorey frugivores (e.g., sedentary birds such as cracids (Cracideae)) feeding on small-fruited plants, are often more restricted in movement than larger-bodied canopy frugivores (e.g., strong-flying hornbills (Bucerotidae)) that favor plants with large fruits (Onstein *et al*., 2017). This segregation may reduce gene flow among plant populations, enhance local adaptation, reproductive isolation, and ultimately speciation (Frankham, 2015, Onstein *et al*., 2017, Méndez *et al*., 2024). Although we did not detect a significant direct relationship between the rate of fruit size evolution and speciation (Fig. 3), this trait may still influence speciation indirectly through correlated evolution with plant height. Moreover, it is plausible that fruit size itself, rather than its rate of evolution, is more important for palm speciation, via frugivory-mediated processes (Onstein *et al*., 2017). Rapid evolutionary changes in fruit size may disrupt palm-frugivore interactions if morphological mismatches arise. This could indicate that speciation in palms may depend more on stabilizing selection exerted by frugivores on maximum fruit sizes given their gape sizes.

Contrary to our hypothesis (H1), we found that slower rates of stem diameter evolution are associated with faster speciation. *Pinanga* and *Ptychosperma* (tribe Areceae), while exhibiting fast leaf size and plant height evolution, showed slow stem diameter evolution (Figs. S5-S7, Table S2). Although stem diameter is crucial for mechanical stability and nutrient/water transport (Givnish, 1995, Laughlin, 2014), palms lack secondary growth, and stem diameter is determined by apical meristem size during early ontogeny (Tomlinson, 2006, Avalos *et al*., 2019). These developmental constraints likely limit the evolutionary trajectory of stem diameter, resulting in lower flexibility of the trait among related lineages. Notably, the negative relationship between stem diameter evolution and speciation was not supported by PGLS analyses, indicating that it may not be robust against phylogenetic autocorrelation, and requires testing across more plant lineages.

### Allometric constraints shape trait evolution and speciation in palms

Our findings in palms supports the ‘allometric constraint hypothesis’ (H2) (Fig. 3). Constraints in palms arise from their unique anatomy, including lack of secondary growth and reliance on a single meristem complex, as noted above (Tomlinson, 2006). This structural and developmental rigidity limits the potential functional spectrum of trait combinations (Díaz *et al*., 2016), restricting phenotypic evolution. For example, rainforest canopy palms maintain allometric relationships between leaf size and plant height to optimize light capture (Göldel *et al*., 2015). Similarly, coordination between stem diameter and plant height ensures mechanical and hydraulic stability (Rich *et al*., 1986, Tomlinson, 2006, Tomlinson & Huggett, 2012). A notable example of trait integration is found in *Geonoma*, where leaf size, stem diameter, and plant height exhibit a strong positive association (Chazdon, 1991). Although palms have extensively explored the structural potential of their arborescent monocot architecture (Tomlinson, 2006), their evolutionary trajectories may have been constrained by anatomical and physiological trade-offs. These limitations may reduce their capacity to fully exploit ecological opportunities, likely restricting speciation. Nevertheless, correlated trait evolution may also open up new ecological opportunities by leading to the development of new trait ‘syndromes’ that could facilitate ecological speciation under divergent selection (Henderson, 2002).

### The complex role of genome size in palm speciation

Genome size has been linked to speciation across angiosperms (Knight *et al*., 2005, Simonin & Roddy, 2018, Moeglein *et al*., 2020). In palms, we found a significant positive association of genome size with speciation rates (Fig. 3), even after excluding the four polyploid species (Fig. S10) and restricting analyses to species with larger genomes (Figs. S11, S12). This finding challenges the ‘large genome constraint hypothesis’ (H3) (Knight *et al*., 2005). A possible explanation is that species with large genomes, primarily produced by repeat accumulation in palms, often occupy more restricted geographic ranges, likely owing to the nucleotypic effects associated with genome size (Knight *et al*., 2005, Bureš *et al*., 2024). Restricted range of species with large genomes may result in smaller population sizes where genetic drift, rather than selection, governs genome dynamics (Bureš *et al*., 2024). In these populations, the reduced efficiency of natural selection may facilitate passive accumulation of repetitive DNA (Lynch *et al*., 2011, Bureš *et al*., 2024). Moreover, the predicted reduced effectiveness of repeat elimination via recombination in palm genomes exceeding 5–6 Gbp/1C allows for non-adaptive mutations to transform repeats into sequences that are no longer recognizable as repeats and hence may evade excision via recombination, further driving genome expansion (Novák *et al*., 2020, Schley *et al*., 2022). Over time, these processes could lead to genetic divergence and reproductive isolation promoting speciation.

Genome size can also impact speciation indirectly through its nucleotypic effect (Bennett, 1971) on morphological traits (Gregory *et al*., 2000, Roddy *et al*., 2020, Carta *et al*., 2022, Feng *et al*., 2022). These genome size-trait correlations can shape trait evolution conferring adaptive advantages or disadvantages depending on the ecological context (Bhadra *et al*., 2023), thereby indirectly affecting speciation. Consistent with the ‘large genome constraint hypothesis’ (H3), we found that genome size likely promotes speciation by facilitating leaf size evolution, while constraining evolution of plant height and stem diameter (Fig. 3). Larger genomes are associated with larger stomatal (guard cell) size and xylem vessel diameter, reducing hydraulic conductance efficiency and potentially limiting evolution of plant height and stem diameter (Knight & Beaulieu, 2008, Feng *et al*., 2022). These constraints may be disadvantageous in water-limited environments hindering the adaptation and establishment of large-genomed palm species (Knight & Beaulieu, 2008, Feng *et al*., 2022, Schley *et al*., 2022). However, in stable, hot and humid rainforest climates where palms predominantly grow, large stomata may not be disadvantageous. In these conditions, reduced environmental selective pressure to minimize water loss may allow plant species with a greater diversity of stomatal sizes to thrive, promoting leaf size evolution and fostering speciation (Veselý *et al*., 2011, Šmarda *et al*., 2023). These findings underscore the complex influence of genome size on speciation of palms mediated by constraints of trait evolution and environmental context.

### Conclusion

Taken together, our study provides comprehensive insights into the complex factors shaping palm speciation across macroevolutionary scales. While environmental pressures play a crucial role, our findings highlight that speciation is related to and constrained by the allometric evolvability of traits and genome size. Specifically, our results are consistent with a scenario where trait-specific evolutionary dynamics and allometric constraints simultaneously facilitate and limit ecological opportunities that drive speciation, in addition to potential effects of genetic drift, rather than selection, on speciation via genome size. By integrating genome size, trait evolution and speciation into a unified framework, we capture the interplay between multiple evolutionary processes. This integrated perspective helps explain how multiple evolutionary processes and traits interact, and how their individual and combined effects shape speciation. Although our focus was on palms, the patterns we identified not only advance our knowledge of this tropical plant family, but also offer a model for investigating similar mechanisms in other (plant) lineages. Ultimately, our work contributes to a deeper understanding of biodiversity and evolutionary dynamics in rainforest ecosystems, and potentially across the plant Tree of Life. Future studies will benefit from development of phylogenetic comparative methods capable of simultaneously disentangling the roles of diverse traits (e.g., functional traits, genome size) and processes (e.g., trait evolution, speciation) (von Hardenberg & Gonzalez-Voyer, 2025), essential for unraveling the drivers of biodiversity.

## Acknowledgments

The work was supported by the German Research Foundation (DFG FZT 118, 202548816), specifically, a personal postdoctoral grant to S.B. (Bhadra) from the Synthesis Centre of iDiv (sDiv). We thank sDiv and Evolution & Adaptation group members of iDiv for their useful discussions on the topic. The authors also thank Gabriele Rada of Media and Communication of iDiv for assisting in preparation of the SEM figure in the manuscript.

## Competing interests

The authors declare no competing interests.

## Author Contributions

S.B. (Bhadra), I.J.L., S.B. (Bellot), W.J.B. and R.E.O. contributed to conceptualizing the project and participated in data compilation. S.B. (Bhadra), I.J.L. and R.E.O. contributed to study design, funding acquisition, and led interpretation of results and conceptual framing. Data analyses were performed by S.B. (Bhadra) and R.E.O. S.B. (Bhadra) coordinated the project, wrote the original draft, and prepared the figures. All authors reviewed and edited the manuscript, and approved the final manuscript.

## Data availability

The data that supports the findings of this study are available in the supplementary material of this article.

## Supplementary information

- **Fig. S1** Pairwise correlations within fruit size and leaf size trait data
- **Fig. S2** Overview of correlations among speciation rates using different approaches
- **Fig. S3** Distribution of prior and posterior, and tip rate comparison from different rate shift analyses in BAMM (Bayesian Analysis of Macroevolutionary Mixtures)
- **Fig. S4** The *a priori* model tested in all the structural equation modeling (SEM) analyses
- **Fig. S5** Palm phylogeny showing rates of speciation and trait evolution
- **Fig. S6** Structural equation model with speciation rates derived from BAMM (Bayesian Analysis of Macroevolutionary Mixtures)
- **Fig. S7** Structural equation model with speciation rates derived from ClaDS (Cladogenetic Diversification rate Shift)
- **Fig. S8** Structural equation model with speciation rates derived from DR (Diversification Rate statistics)
- **Fig. S9** Structural equation model with speciation rates estimated from BAMM (Bayesian Analysis of Macroevolutionary Mixtures) illustrating the variations in coefficient values when accounted for phylogenetic tree topology
- **Fig. S10** Structural equation model excluding the polyploids (outliers)
- **Fig. S11** Structural equation model including only species with genome size higher than the median genome size value (i.e., > 2.64 Gbp/1C)
- **Fig. S12** Structural equation model including species above the 75th percentile range of genome size (i.e., > 4.26 Gbp/1C)
- **Table S1** Palm trait and genome size data used in the analyses
- **Table S2** Compiled dataset of the estimated rates of speciation and trait evolution of each species
- **Table S3** Data of prior information used for BAMM (Bayesian Analysis for Macroevolutionary Mixtures) analysis
- **Table S4** Results from the phylogenetic generalized least squares (PGLS) analyses
- **Notes S1** Addressing criticism against BAMM

## Notes

### Competing Interest Statement

The authors have declared no competing interest.

