## Supplementary information for "Functional traits drive speciation in tropical palms through complex interactions between genome size, adaptation and allometry"

### Supplementary figures

**Fig. S1 Pairwise correlations within fruit size and leaf size trait data.** The figure shows the correlations among the trait categories of (a) fruit size and (b) leaf size. The upper diagonal contains the values of the Pearson's correlation coefficient (r) with the asterisk (*) indicating significant p-value (p<0.05). The lower diagonal contains the scatterplots with the x- and y- axis indicating the log-transformed trait data. The trend line (in red) in the scatter plots indicate the linearity of the positive relationship between the trait data.

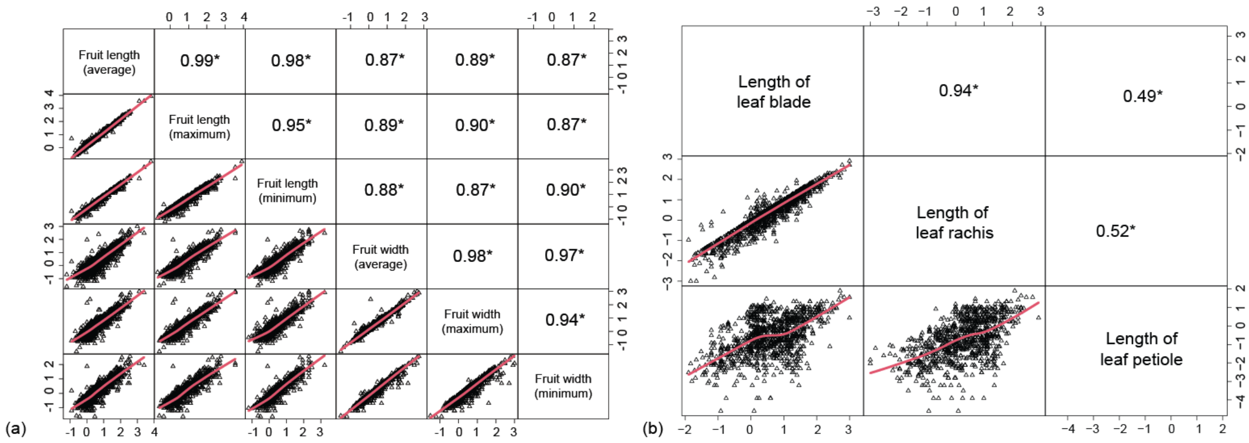

**Fig. S2 Overview of correlations among speciation rates using different approaches.** Pairwise correlations among the tip rates of speciation derived from BAMM (Bayesian Analysis of Macroevolutionary Mixtures), ClaDS (Cladogenetic Diversification rate Shift) and DR (Diversification Rate statistics) in lineages per million years. The upper diagonal contains the values of the Pearson's correlation coefficient (r) with the asterisk (*) indicating significant p-value ($p<0.05$). The lower diagonal contains the scatterplots with the x- and y- axis indicating tip rates. The trend line (in red) in the scatter plots indicate the positive linearity of relationships between the tip rates.

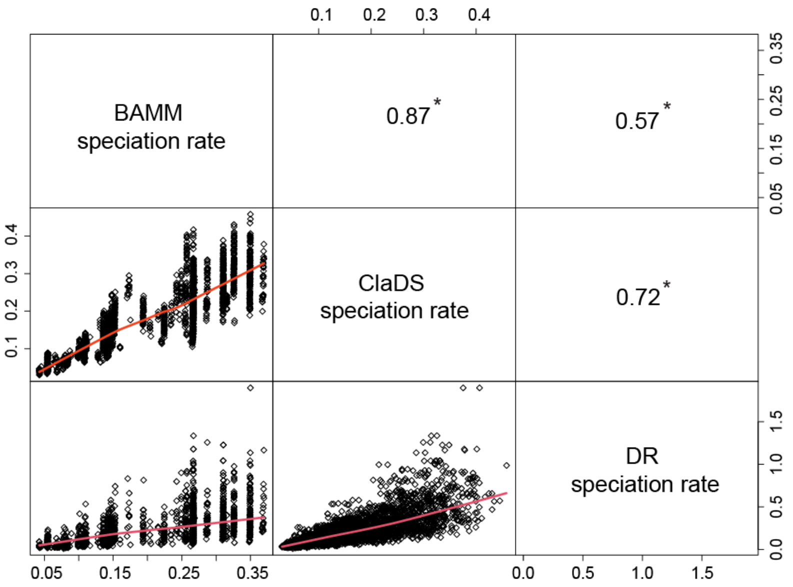

**Fig. S3 Distribution of prior and posterior, and tip rate comparison from different rate shift analyses in BAMM (Bayesian Analysis of Macroevolutionary Mixtures).** (a-c) Distribution of prior and posterior in BAMM analyses using different prior values (i.e., expected number of shifts): (a) 1, (b) 5, and (c) 10. The x-axis represents the number of rate shifts and the y-axis represents the frequency of rate shifts observed in the prior or posterior distribution represented by blue and yellow histogram respectively. The prior and posterior distributions are distinct and non-overlapping in all cases. With increasing values of the prior on the expected number of shifts, the prior distribution becomes less informative, and the posterior distribution becomes wider. (d) Comparison of tip-rates derived from the three different shift configurations. Upper diagonal contains the values of Pearson's correlation coefficient (r). The asterisk (*) indicates $p<0.05$. All the shifts showed almost similar tip rates as indicated by the Pearson's r values. The lower diagonal contains the scatterplots with the x- and y- axis indicating tip rates. The trend line (in red) in the scatter plots indicate the positive and linear relationship between the tip rates.

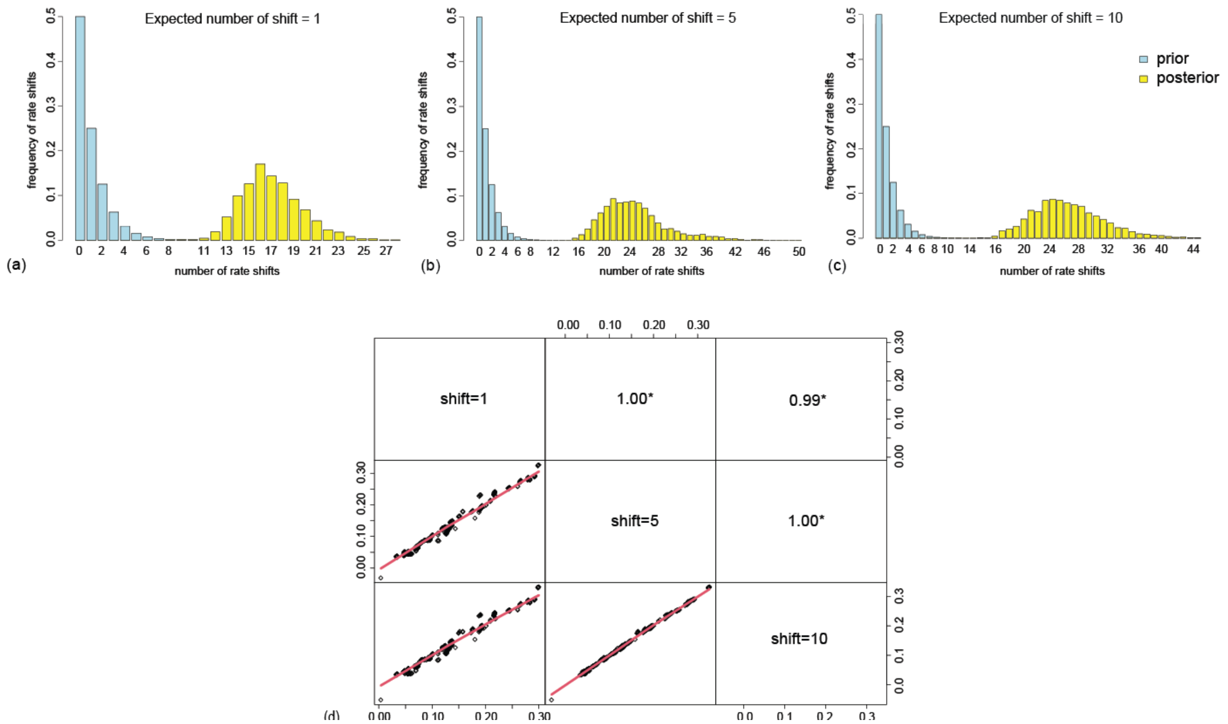

**Fig. S4 The *a priori* model tested in all the structural equation modeling (SEM) analyses.** The figure depicts the initial model tested in all SEM analyses. It includes the effects of trait evolutionary rates of plant height, stem diameter, leaf size and fruit size on speciation rates (trait flexibility hypothesis, H1), correlations between trait evolution rates (allometric constraint hypothesis, H2), and of genome size on speciation (genome size constraint hypothesis, H3). It shows all the hypothetical pathways based on the information obtained by testing correlations between each variable and theoretical considerations. The black arrows represent hypothesized positive relationships and the red arrows represent the hypothesized negative relationships.

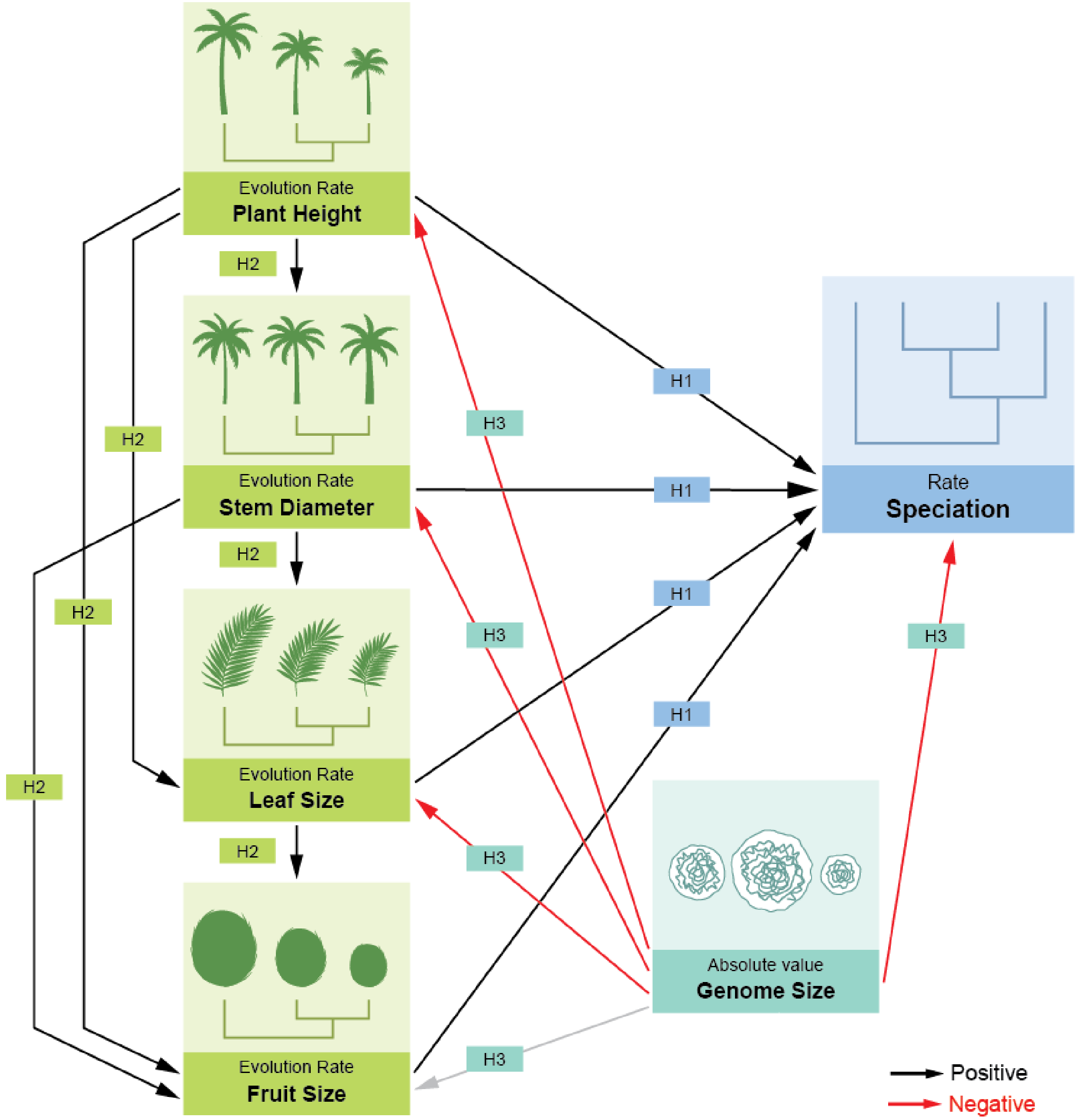

**Fig. S5 Palm phylogeny showing rates of speciation and trait evolution.** Phylogenetic trees showing BAMM-derived estimates of (a) speciation rate, (b) plant height evolution rate, (c) stem size evolution rate, (d) leaf size evolution rate, and (e) fruit size evolution rate. Black vertical bars on the right side of each phylogeny indicate lineages exhibiting evolutionary radiations (i.e., speciation rates estimated to be above background speciation rates). Vertical coloured bars indicate palm subfamilies. Detailed subfamily boundaries and species names are visible upon zooming into the figure.

**Fig. S6 Structural equation model with speciation rates derived from BAMM (Bayesian Analysis of Macroevolutionary Mixtures).** The figure shows the effects of trait evolutionary rates of plant height, stem diameter, leaf size and fruit size on speciation rates (trait flexibility hypothesis, H1), correlations between trait evolution rates (allometric constraint hypothesis, H2), and of genome size on speciation (genome size constraint hypothesis, H3) in palms (N=372 species) with speciation rate derived from BAMM (Bayesian Analysis of Macroevolutionary Mixtures). The effect sizes indicate standardized coefficients with significance ($p<0.05$). The arrow thickness is proportional to coefficient values and the arrow direction represents the direction of effects. Black arrows denote positive effects, red arrows denote negative effects, and grey arrows denote tested but statistically unsupported effects ($p>0.05$). Fit indices of the model were as follows: p-value= 0.525, CFI= 1.000, TLI= 1.017, RMSEA= 0.000, SRMR= 0.014.

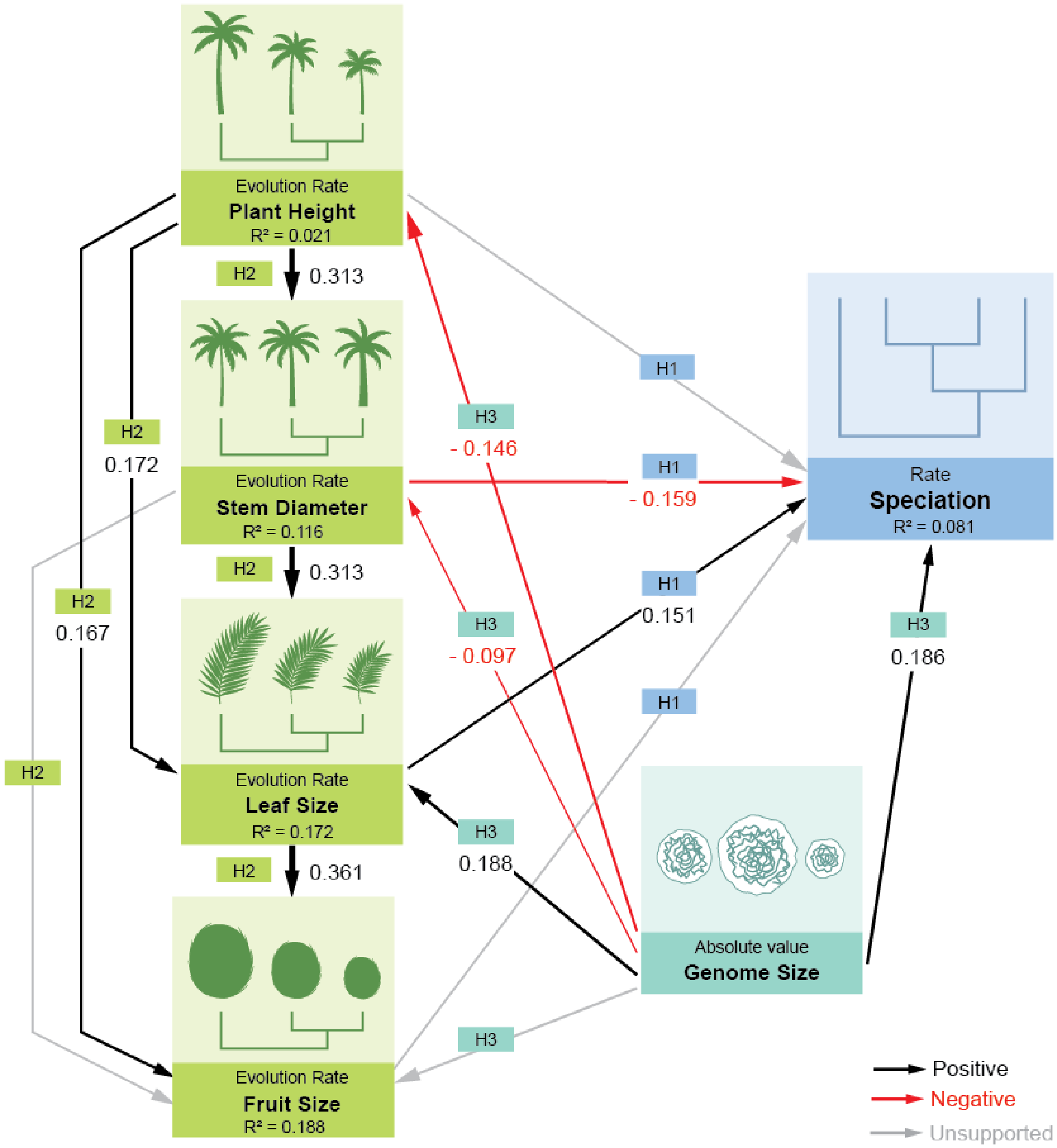

**Fig. S7 Structural equation model with speciation rates derived from ClaDS (Cladogenetic Diversification rate Shift).** The figure shows the effects of trait evolutionary rates of plant height, stem diameter, leaf size and fruit size on speciation rates (trait flexibility hypothesis, H1), correlations between trait evolution rates (allometric constraint hypothesis, H2), and of genome size on speciation (genome size constraint hypothesis, H3) in palms (N=372 species) with speciation rate derived from ClaDS (Cladogenetic Diversification rate Shift). The effect sizes indicate standardized coefficients with significance ($p<0.05$). The arrow thickness is proportional to coefficient values and the arrow direction represents the direction of effects. Black arrows denote positive effects, red arrows denote negative effects, and grey arrows denote tested but statistically unsupported effects ($p>0.05$). Fit indices of the model were as follows: p-value= 0.667, CFI= 1.000, TLI= 1.038, RMSEA= 0.000, SRMR= 0.008.

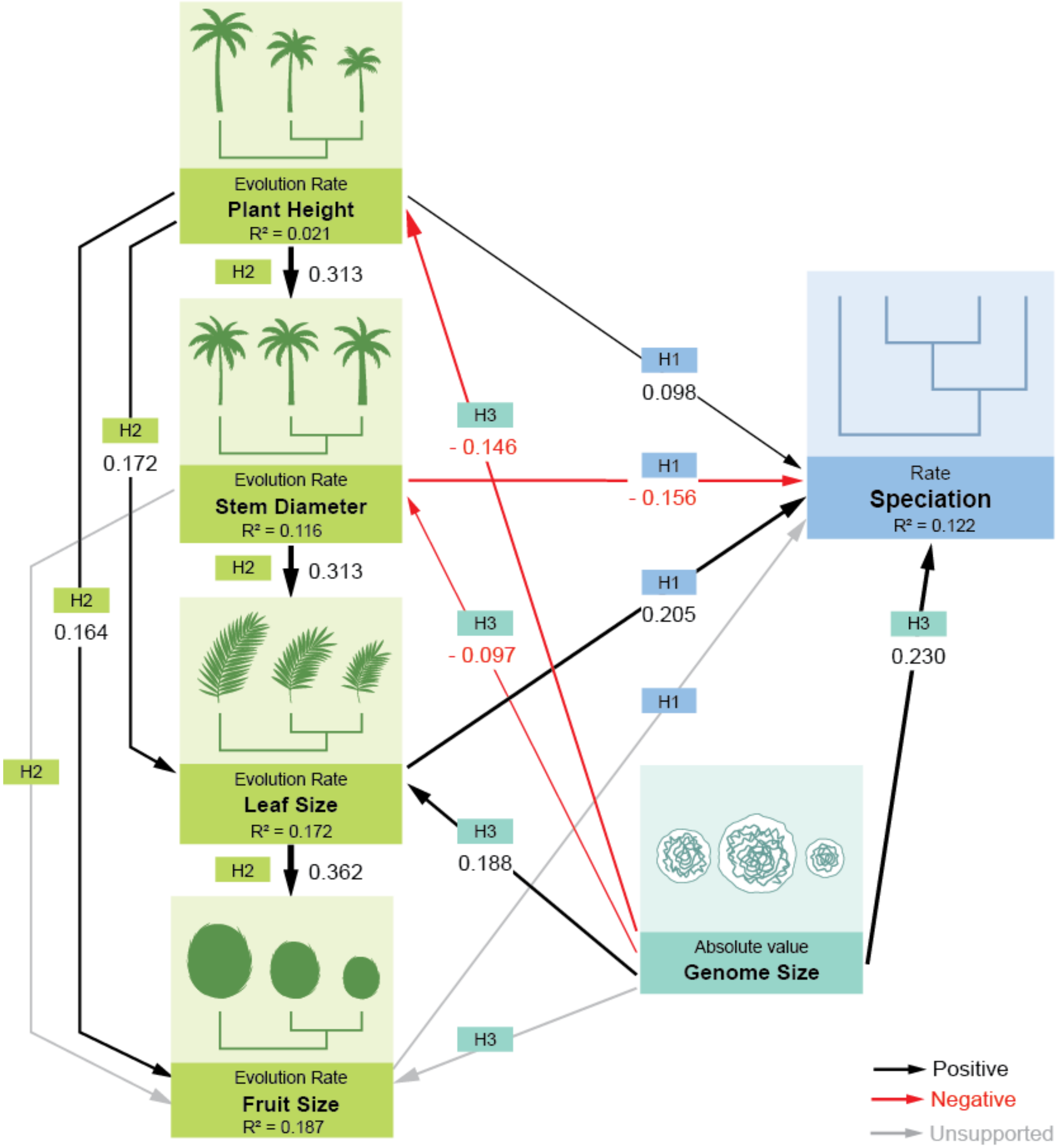

**Fig. S8 Structural equation model with speciation rates derived from DR (Diversification Rate statistics).** The figure shows the effects of trait evolutionary rates of plant height, stem diameter, leaf size and fruit size on speciation rates (trait flexibility hypothesis, H1), correlations between trait evolution rates (allometric constraint hypothesis, H2), and of genome size on speciation (genome size constraint hypothesis, H3) in palms (N=372 species) with speciation rate derived from DR (Diversification rate statistics). The effect sizes indicate standardized coefficients with significance ($p<0.05$). The arrow thickness is proportional to coefficient values and the arrow direction represents the direction of effects. Black arrows denote positive effects, red arrows denote negative effects, and grey arrows denote tested but statistically unsupported effects ($p>0.05$). Fit indices of the model were as follows: p-value= 0.595, CFI= 1.000, TLI= 1.024, RMSEA= 0.000, SRMR= 0.013.

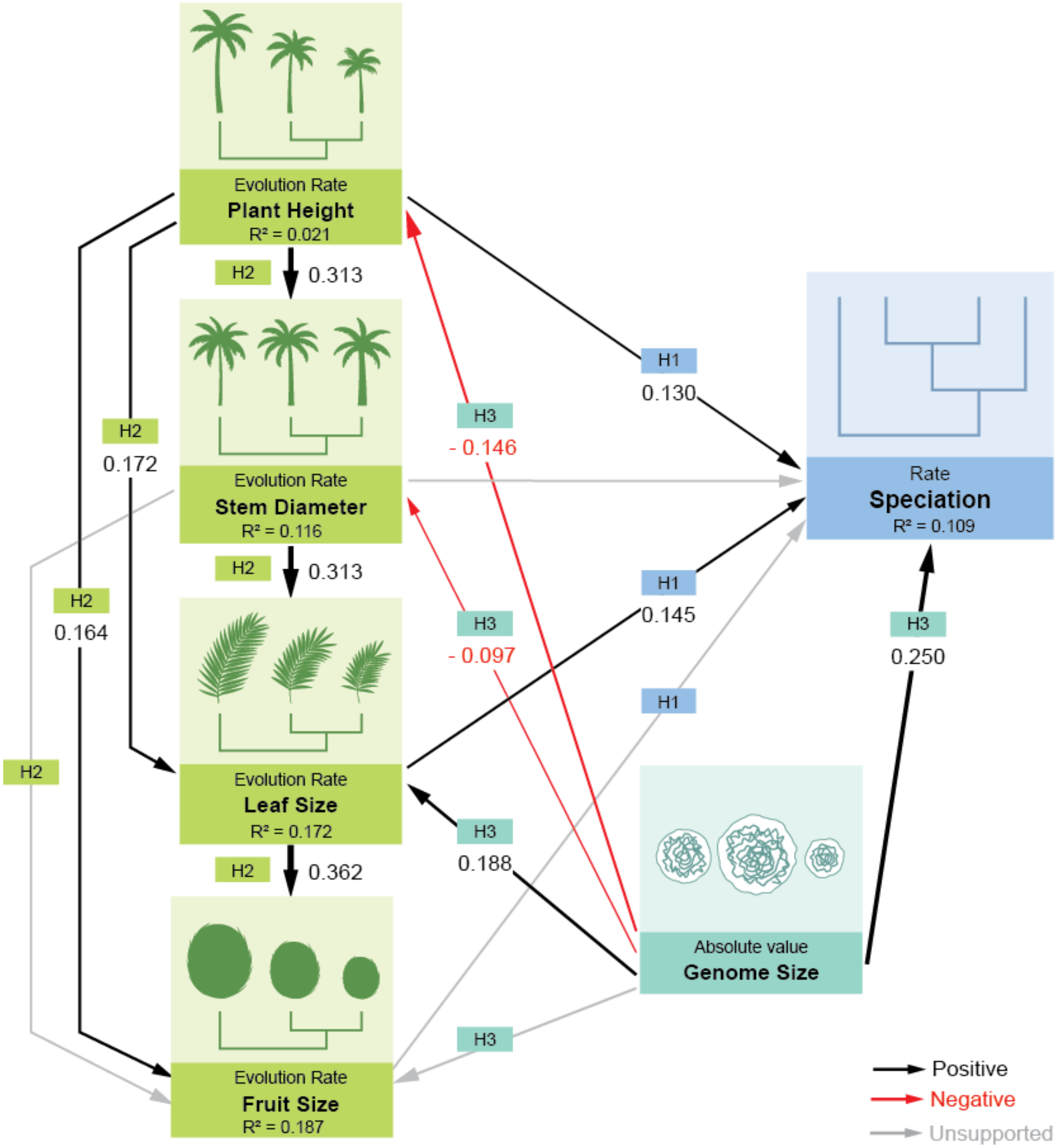

**Fig. S9 Structural equation model with speciation rates estimated from BAMM (Bayesian Analysis of Macroevolutionary Mixtures) illustrating the variations in coefficient values when accounted for phylogenetic tree topology.** The figure presents the final structural equation model (SEM) using BAMM-derived speciation rates from the MCC (Maximum Clade Credibility) phylogenetic tree. The hypotheses tested are the effects of trait evolutionary rates of plant height, stem diameter, leaf size and fruit size on speciation rates (trait flexibility hypothesis, H1), correlations between trait evolution rates (allometric constraint hypothesis, H2), and of genome size on speciation (genome size constraint hypothesis, H3). Alphabets on the arrows (a-e) in the SEM diagram indicate relationships for which standardized coefficient values varied when SEMs were performed across 100 constrained, posterior-distribution phylogenetic trees of palms. The plots (a-e) show the distribution of standardized coefficients for these relationships, highlighting where they differed from the SEM values obtained using the MCC tree. The plots include correlations between (a) genome size and speciation rate, (b) stem diameter evolution rate and speciation rate, (c) leaf size evolution rate and speciation rate, (d) leaf size evolution rate and fruit size evolution rate, and (e) plant height evolution rate and leaf size evolution rate. In each plot, the x-axis indicates the standardized coefficient values, the y-axis shows their frequency across the 100 trees, blue lines represent the 5th and 95th quantiles of the coefficients, and the red line marks the coefficient value obtained from the SEM based on the MCC tree.

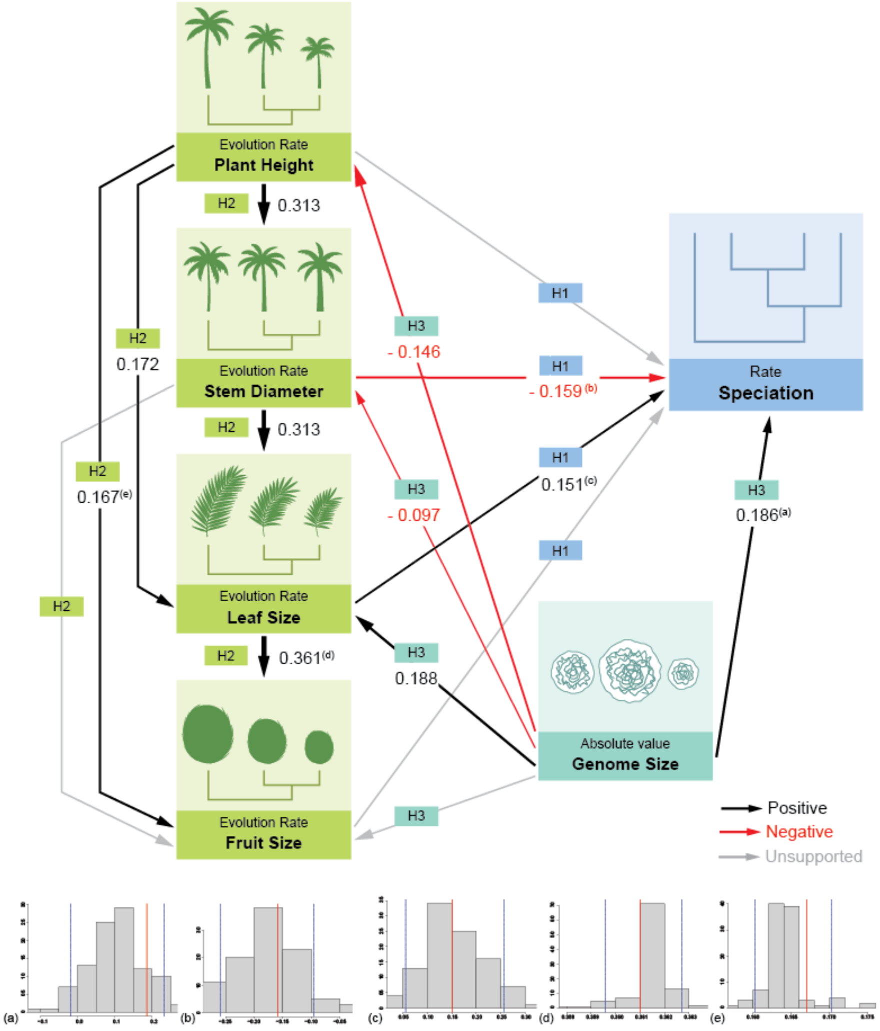

**Fig. S10 Structural equation model excluding the polyploids (outliers).** The figure shows the effects of trait evolutionary rates of plant height, stem diameter, leaf size and fruit size on speciation rates (trait flexibility hypothesis, H1), correlations between trait evolution rates (allometric constraint hypothesis, H2), and of genome size on speciation (genome size constraint hypothesis, H3) in palms with speciation rate derived from BAMM (Bayesian Analysis of Macroevolutionary Mixtures). The analysis excludes the polyploid species (N=4) which were considered as outliers in the study. The effect sizes indicate standardized coefficients with significance ($p<0.05$). The arrow thickness is proportional to coefficient values and the arrow direction represents the direction of effects. Black arrows denote positive effects, red arrows denote negative effects, and grey arrows denote tested but statistically unsupported effects ($p>0.05$). Fit indices of the model were as follows: p-value= 0.593, CFI= 1.000, TLI= 1.024, RMSEA= 0.000, SRMR= 0.013.

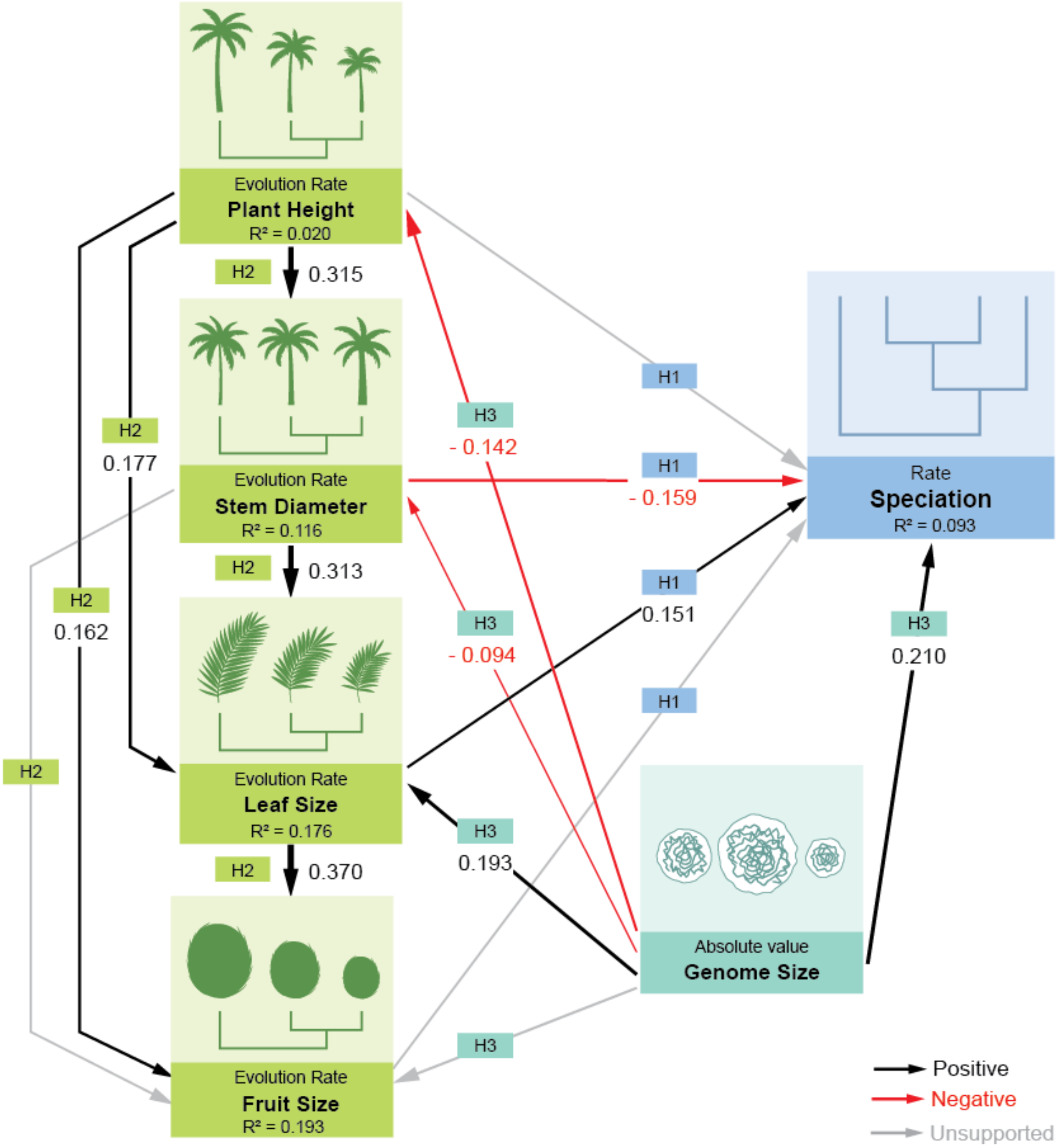

**Fig. S11 Structural equation model including only species with genome size higher than the median genome size value (i.e., > 2.64 Gbp/1C).** The figure shows the effects of trait evolutionary rates of plant height, stem diameter, leaf size and fruit size on speciation rates (trait flexibility hypothesis, H1), correlations between trait evolution rates (allometric constraint hypothesis, H2), and of genome size on speciation (genome size constraint hypothesis, H3) in palms (N=189 species) with speciation rate derived from BAMM (Bayesian Analysis of Macroevolutionary Mixtures). Species that possess genome size higher than the median genome size of palms (i.e., 1C > 2.64 Gbp) are included. The effect sizes indicate standardized coefficients with significance ($p<0.05$). The arrow thickness is proportional to coefficient values and the arrow direction represents the direction of effects. Black arrows denote positive effects, red arrows denote negative effects, and grey arrows denote tested but statistically unsupported effects ($p>0.05$). Fit indices of the model were as follows: p-value= 0.943, CFI= 1.000, TLI= 1.061, RMSEA= 0.000, SRMR= 0.019.

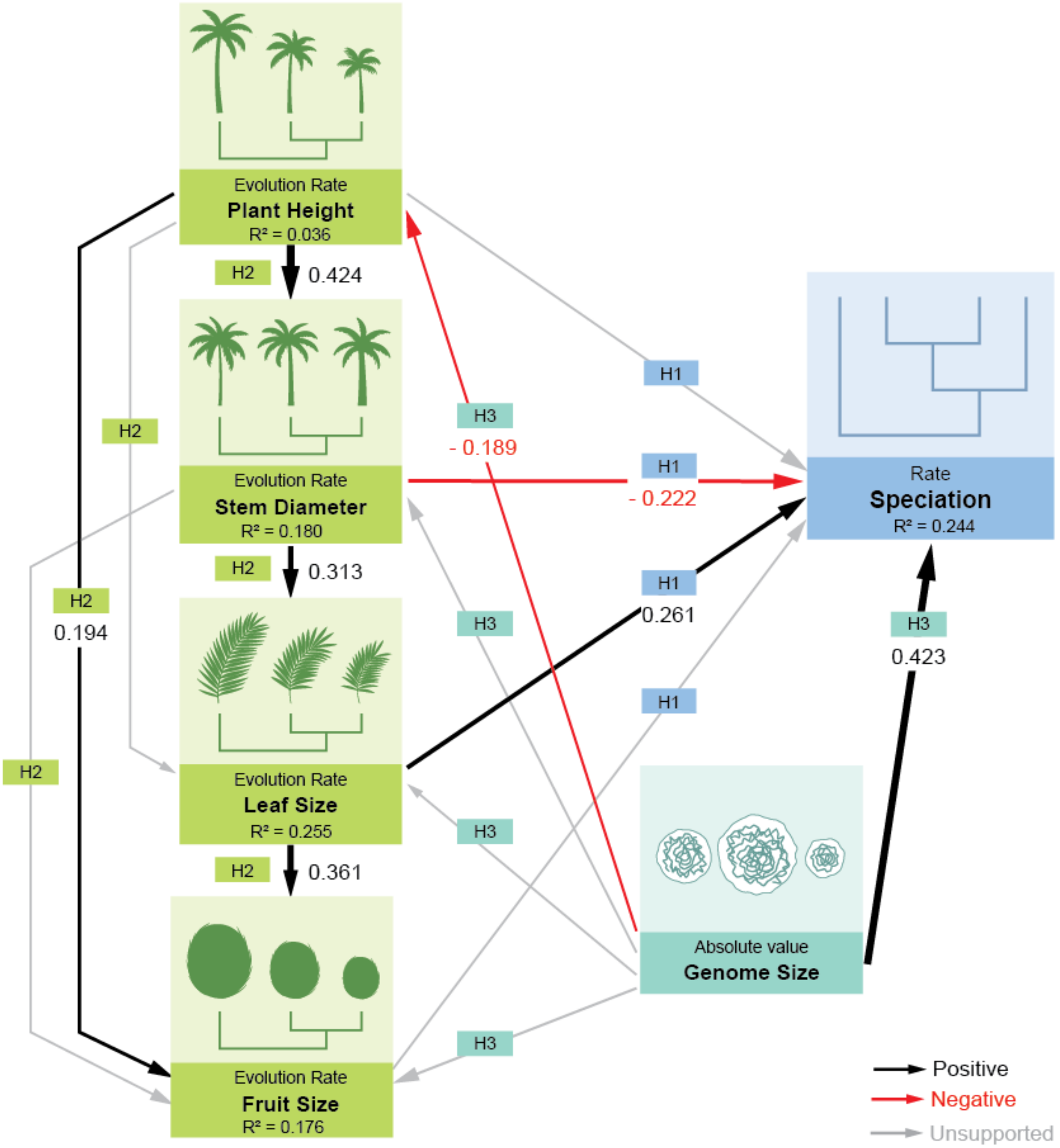

**Fig. S12 Structural equation model including species above the 75th percentile range of genome size (i.e., > 4.26 Gbp/1C).** The figure shows the effects of trait evolutionary rates of plant height, stem diameter, leaf size and fruit size on speciation rates (trait flexibility hypothesis, H1), correlations between trait evolution rates (allometric constraint hypothesis, H2), and of genome size on speciation (genome size constraint hypothesis, H3) in palms (N=95 species) with speciation rate derived from BAMM (Bayesian Analysis of Macroevolutionary Mixtures). Species that possess genome size above the 75th percentile range of genome size of palms (i.e., 1C > 4.26 Gbp) are included. The effect sizes indicate standardized coefficients with significance ($p<0.05$). The arrow thickness is proportional to coefficient values and the arrow direction represents the direction of effects. Black arrows denote positive effects, red arrows denote negative effects, and grey arrows denote tested but statistically unsupported effects ($p>0.05$). Fit indices of the model were as follows: p-value= 0.407, CFI= 0.998, TLI= 0.996, RMSEA= 0.016, SRMR= 0.049.

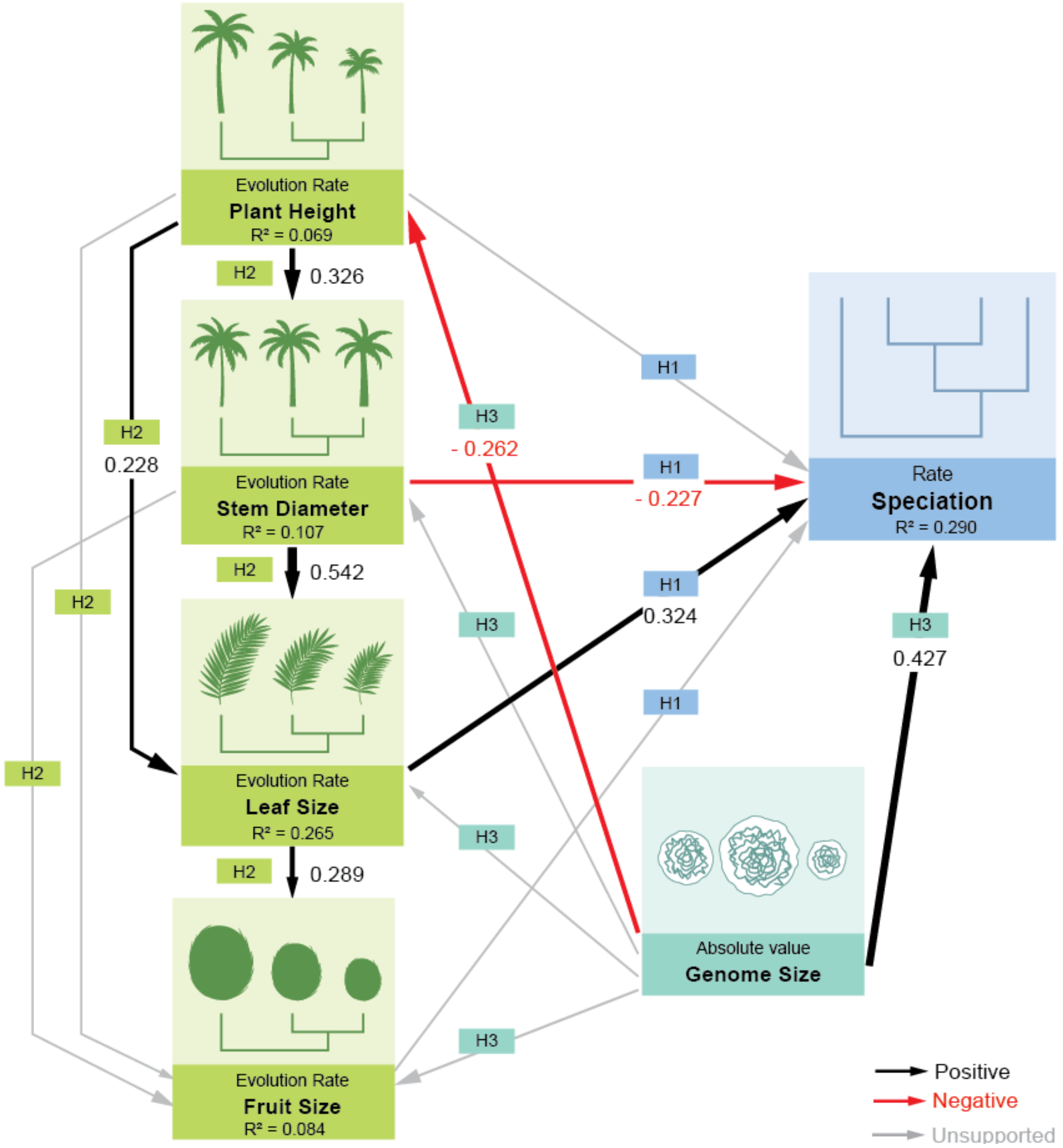

### Supplementary Tables

**Table S1 Palm trait and genome size data used in the analyses.** The table lists all traits and genome size data included in the initial analyses. The number and proportion of species for which data were available is also included. Traits shown in bold were included in the final analysis.

| Plant traits | Name of the variable | Total number of species with data (%) |
|---|---|---|
| | Total number of species | 2528 |
| Genome size | **Genome size** | **425 (16.71)** |
| Fruits | **Average fruit length** | **2041 (80.23)** |
| | Maximum fruit length | 911 (35.81) |
| | Minimum fruit length | 901 (35.42) |
| | Average fruit width | 1983 (77.95) |
| | Maximum fruit width | 997 (39.19) |
| | Minimum fruit width | 989 (38.87) |
| Stem | **Maximum plant height** | **1985 (78.03)** |
| | **Maximum stem diameter** | **1929 (75.82)** |
| Leaf | **Maximum leaf blade length** | **1892 (74.37)** |
| | Maximum leaf rachis length | 1524 (59.91) |
| | Maximum leaf petiole length | 1202 (47.25) |

**Table S2 Compiled dataset of the estimated rates of speciation and trait evolution of each species.** This table presents a detailed dataset derived from our analyses, including estimates of speciation rate (in lineages per million years) using three methods: BAMM (Bayesian Analysis of Macroevolutionary Mixtures), ClaDS (Cladogenetic Diversification rate Shift) and DR (Diversification Rate statistics), measured from the MCC phylogenetic tree (Faurby et al., 2016). Trait evolution rates (in lineages per million years) derived from the MCC phylogenetic tree (Faurby *et al*., 2016) is included. Genome size data (in Gbp/1C) were compiled from previously published resources (Pellicer & Leitch, 2020, Schley *et al*., 2022). Columns labelled BAMM1 to BAMM100 represent speciation rates measured from 100 constrained, posterior phylogenetic trees (Faurby *et al*., 2016) using BAMM. For consistency, nomenclature of Faurby *et al*. (2016) was used in our analyses. For taxonomic clarity, the accepted species names according to the World Checklist of Vascular Plants (WCVP) are also included, along with their corresponding taxonomic status. The higher order classification (subfamily, tribe and genus) is also included.

**Table S3 Data of prior information used for BAMM (Bayesian Analysis for Macroevolutionary Mixtures) analysis.** The data was generated using the setBAMMPriors() function of the 'BAMMtools' package in R, which was applied to analyze speciation-extinction rate and trait evolution rate in BAMM.

| Analyses | Expected number of shifts | Lambda initial prior | Lambda shift prior | Mu initial prior | Beta initial prior | Beta shift prior |
|---|---|---|---|---|---|---|
| Speciation-extinction | 1 | 2.9511 | 0.0109 | 2.9511 | – | – |
| Phenotypic evolution | | | | | | |
| — Plant height | 1 | – | – | – | 0.0108 | 0.0109 |
| — Stem diameter | 1 | – | – | – | 0.0091 | 0.0109 |
| — Leaf size | 1 | – | – | – | 0.3688 | 0.0109 |
| — Fruit size | 1 | – | – | – | 0.4785 | 0.0109 |

**Table S4 Results from the phylogenetic generalized least squares (PGLS) analyses.** The table reports results of PGLS assessing phylogenetic autocorrelation between response and predictor variables included in the final structural equation models (SEM) using BAMM-, ClaDS- and DR-derived speciation rates. For each model, the estimated phylogenetic signal (λ) is reported (N= 372). For each predictor variable, the parameter estimate (β), standard error (SE), t-statistic and p-value are provided. Statistically significant relationships ($p < 0.05$) are marked with an asterisks (*).

| Response variable (Speciation rates) | Predictor variables | Variable statistics | | | | Model statistics |
|---|---|---|---|---|---|---|
| | | β | SE | t | p-value | λ |
| BAMM | Genome size | -0.003 | 0.01 | -0.22 | 0.82 | 1.00 |
| | Leaf size evolution rate | 0.035 | 0.03 | 1.29 | 0.19 | |
| | Stem diameter evolution rate | -0.017 | 0.03 | -0.63 | 0.52 | |
| ClaDS | Genome size | 0.014 | 0.04 | 0.37 | 0.71 | 0.998 |
| | Leaf size evolution rate | 0.073 | 0.03 | 2.43 | 0.01* | |
| | Stem diameter evolution rate | 0.004 | 0.03 | 0.14 | 0.89 | |
| | Plant height evolution rate | 0.091 | 0.03 | 2.81 | 0.005* | |
| DR | Genome size | 0.013 | 0.02 | 0.51 | 0.61 | 1.00 |
| | Leaf size evolution rate | 0.196 | 0.05 | 4.17 | 3.816e-05* | |
| | Plant height evolution rate | 0.126 | 0.05 | 2.48 | 0.0135* | |

### Supplementary Notes

**Notes S1** Addressing criticism against BAMM

The macroevolutionary dynamics of speciation and trait evolution of palms were investigated using the C++ programme BAMM v. 2.5.0 (Rabosky, 2014). BAMM uses Bayesian model to estimate the discrete rate shifts for each node, and its descendants, to assess the diversification rates (speciation-extinction parameter) and trait evolution rates (phenotypic evolution parameter). It uses reversible-jump Markov Chain Monte Carlo (rjMCMC) that implements a compound Poisson process (CPP). CPP allows for heterogeneity and random distribution of rate shifts along the branches of a phylogenetic tree and through evolutionary time (Rabosky, 2014).

The BAMM framework facilitates the simulation of posterior distribution of shift configurations. Each shift configuration corresponds to a set of evolutionary "shifts" which represents variations in rates of speciation and extinction along specific branches of a phylogenetic tree, and subsequent mapping of these shifts across the entire phylogenetic tree (Rabosky, 2014, Rabosky *et al*., 2017). The expected complexity of the speciation process of the phylogeny is set by the anticipated number of rate shifts (expectedNumberOfShifts parameter) in the tree estimated based on prior information and the model used. Prior presumes the distribution of shifts to be uniform, and the specified number of shifts together with the branch length influences the probability of finding the posterior distribution of a shift (Rabosky, 2014, Rabosky *et al*., 2017). Higher values of expected shifts in prior will result in higher shifts in posterior (Rabosky, 2014, Rabosky *et al*., 2017). The resulting posterior distribution allows to generate a "phylorate" plot that facilitates the visual representation of the macroevolutionary dynamics of speciation and trait evolution rates. The phylorate plot is constructed by averaging and plotting all the shift configurations within the posterior distribution on each time unit for each branch (Rabosky, 2014, Rabosky *et al*., 2017).

There is controversy regarding the estimation of rate shifts and calculation of tip rates (i.e., recent rates; measured as lineages per million years) for speciation and trait evolution across a phylogenetic tree by BAMM, primarily due to the sensitivity of posterior distribution to assumed prior shifts and non-inclusion of rate shifts in extinct branches of a tree (Moore *et al*., 2016, Meyer *et al*., 2018). Rabosky et al. (2017) advocated the robustness of BAMM, and subsequent studies have proved that the estimates of speciation patterns by BAMM to be realistic (Magallón *et al*., 2018, Rabosky, 2020, Martínez-Gómez *et al*., 2023). In the present study, our primary focus was the tip rates, and not the location and number of rate shifts (i.e., historical bursts of speciation) that occurred in particular lineages. Hence, we do not have grounds to suspect potential bias in the estimates of the rates of speciation and trait evolution.

However, recognizing the potential problems, we exercised caution while using the tip rates of BAMM for our study and assessed whether the sensitivity of prior on posterior distribution affects the tip rates in our analyses. BAMM provides the setBAMMpriors function in the 'BAMMtools' package in R (Rabosky *et al*., 2014) that is used to scale the priors on rate parameters to the analyzed phylogenetic tree and is also used for post-run analysis. Using the results of setBAMMpriors (Table S3), the rate parameter of exponential prior for the initial speciation (lambdaInitPrior) and extinction (muInitPrior) values were both set to 2.9511. The prior for standard deviation of the normal distribution (mean fixed at zero) of speciation shift parameter for rate regimes was set to 0.0109. Rates were allowed to vary through time (lambdaIsTimeVariablePrior = 1). The expectedNumberOfShifts parameter was set to 1, as suggested by the setBAMMpriors parameter. To test the sensitivity of posterior distribution on the prior, we further conducted two independent analyses to estimate speciation rates over a wide range of magnitudes for the prior setting with the expectedNumberOfShifts parameter being set at 5 and 10. Here 1 assumes a lower probability of observing multiple rate shifts in the phylogenetic tree while 10 represents a high probability of observing multiple rate shifts, based on the number of taxa in the analysis. For each shift configuration, MCMC simulation consisted of 4 independent chains of 300 million generations each with shift configuration being sampled every 10,000 steps. Run convergence was assessed by calculating the effective sampling size for the likelihood of the data and for the number of rate shifts in each regime using the *effectiveSize()* function of the 'coda' package (Plummer *et al*., 2006). The initial 10% of MCMC was discarded as burn-in that resulted in a total of 270,001 analyzed posterior samples. The number of MCMC generations and the burn-in that was used to estimate diversification rates were chosen based on the visual inspection of the log-likelihood of the MCMC output (Barreto *et al*., 2024). After removing 10% of trees as burn-in, we analyzed BAMM output using BAMMtools (Rabosky *et al*., 2014). Tip rates of speciation for each shift configuration were extracted using the *getTipRates()* function. We compared the similarity between the tip rates using Pearson's r correlation coefficient in R.

The posterior distribution was distinctly decoupled from the prior in all the rate shift configurations in our analyses (Fig. S3a-c) suggesting the independence of posterior distributions. Further, the tip rates derived from the three shift configurations were highly similar as was indicated by the values of Pearson's r that showed a significant ($p<0.05$) range of 0.99 to 1.00 (Fig. S3d). Hence, we conducted our main analyses with the expectedNumberOfShifts being set to 1, as suggested by BAMM.
