## Supplementary material for "Functional traits drive speciation in tropical palms through complex interactions between genome size, adaptation and allometry": Fig. S5

**Fig. S5 Palm phylogeny showing rates of speciation and trait evolution.** Phylogenetic trees showing BAMM-derived estimates of (a) speciation rate, (b) plant height evolution rate, (c) stem size evolution rate, (d) leaf size evolution rate, and (e) fruit size evolution rate. Black vertical bars on the right side of each phylogeny indicate lineages exhibiting evolutionary radiations (i.e., speciation rates estimated to be above background speciation rates). Vertical coloured bars indicate palm subfamilies. Detailed subfamily boundaries and species names are visible upon zooming into the figure.

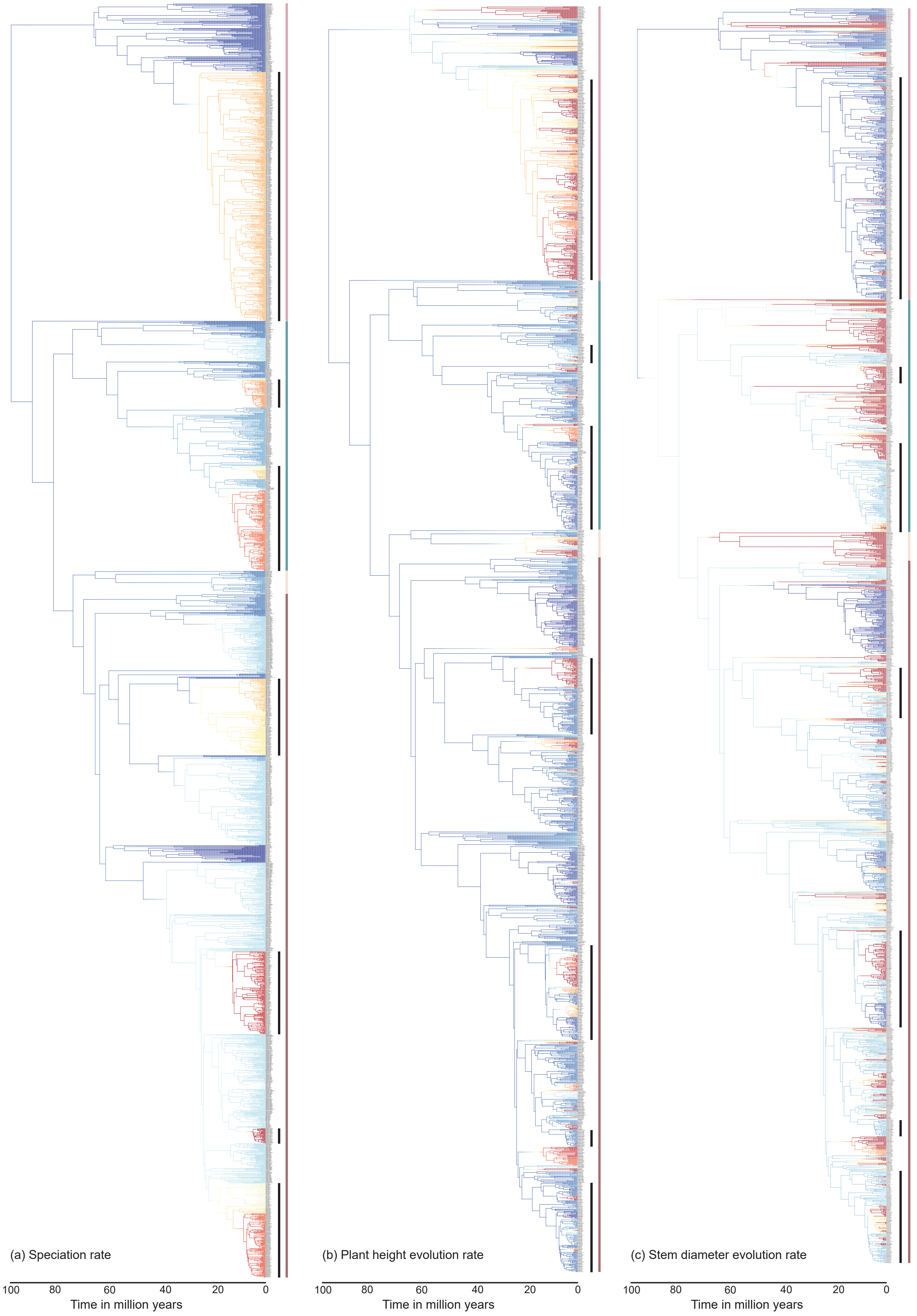

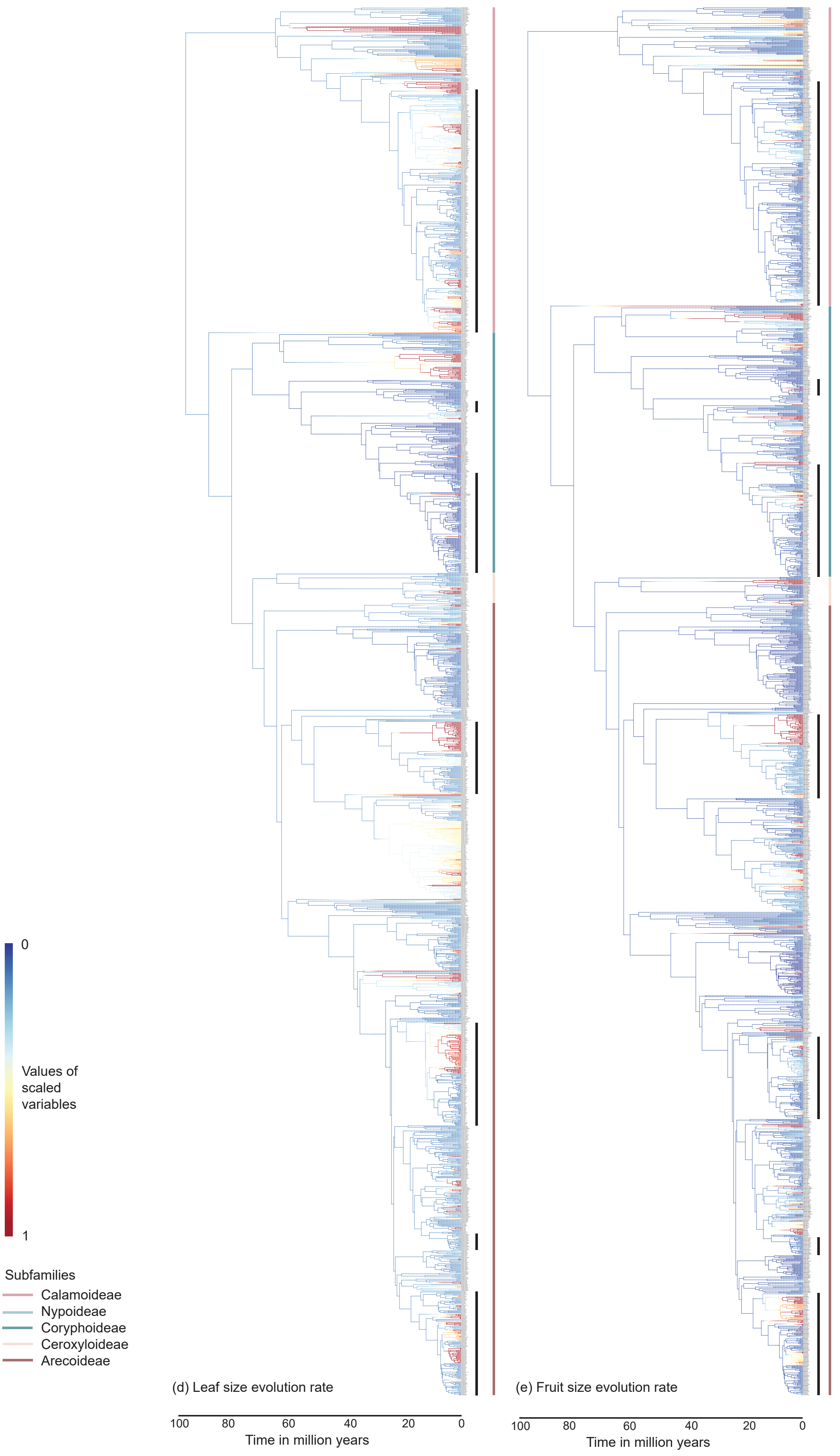

0
Values of scaled variables
1
Subfamilies
Calamoideae
Nypoideae
Coryphoideae
Ceroxyloideae
Arecoideae
(d) Leaf size evolution rate
100
80
60
40
20
0
Time in million years
(e) Fruit size evolution rate
100
80
60
40
20
0
Time in million years
